# An E3 ligase network engages GCN1 to promote elongation factor-1*α* degradation on stalled ribosomes

**DOI:** 10.1101/2022.03.21.485216

**Authors:** Keely Oltion, Jordan D. Carelli, Tangpo Yang, Stephanie K. See, Hao-Yuan Wang, Martin Kampmann, Jack Taunton

**Affiliations:** Chemistry and Chemical Biology Graduate Program, University of California San Francisco, San Francisco, CA, USA; Department of Cellular and Molecular Pharmacology, University of California San Francisco, San Francisco, CA, USA; Institute for Neurodegenerative Diseases, Department of Biochemistry and Biophysics, University of California San Francisco, San Francisco, United States; Chan-Zuckerberg Biohub, San Francisco, CA, USA

## Abstract

How cells monitor the status of translating ribosomes is a major question in gene regulation. Elongating ribosomes frequently stall during mRNA translation, resulting in context-dependent activation of quality control pathways. However, surveillance mechanisms that specifically respond to stalled ribosomes with an elongation factor occupying the GTPase center have not been identified. By employing ternatin-4, an allosteric elongation factor-1*α* (eEF1A) inhibitor, we unveil an E3 ligase network that triggers ubiquitination and degradation of eEF1A on stalled ribosomes. A CRISPRi screen revealed two E3 ligases of unknown function, RNF14 and RNF25, which are both essential for ternatin-induced eEF1A degradation. Based on quantitative proteomics analysis, we find that RNF14 and RNF25 promote ubiquitination of eEF1A and a discrete set of ribosomal proteins. By forming a complex with RNF14, the ribosome collision sensor GCN1 plays an essential role in eEF1A degradation. Our findings illuminate a translation elongation checkpoint that monitors the ribosomal GTPase center.

## Introduction

Protein synthesis is essential for growth and survival in all organisms. Each stage of protein synthesis, including initiation, elongation, and termination, is choreographed by factors that interact with the central catalytic machinery – the ribosome (Dever and Green, 2012). The initiation stage, often rate limiting for protein production, is tightly regulated by cellular signaling pathways (Jackson et al., 2010; Sonenberg and Hinnebusch, 2009). We know less about how cells regulate elongation rates, which are highly variable across coding sequences. Such regulation underlies, for example, the dependence of elongation rates on peptide sequence, codon usage, tRNA expression, and post-transcriptional modifications of tRNA and mRNA (Richter and Coller, 2015; Schuller and Green, 2018). As a consequence, ribosomes may pause or stall in a context-dependent manner, and this may be critical for optimal folding or subcellular targeting of a nascent polypeptide.

Ribosome stalling can also result from attempted translation of a defective mRNA (Brandman and Hegde, 2016; Yip and Shao, 2021). Such pathological stalls can lead to proteotoxic stress caused by depletion of active ribosomes and accumulation of partially synthesized nascent polypeptides. Mechanistic studies using mRNA reporters with poly(A) and other stall-prone coding sequences have revealed surveillance pathways that recognize distinct structural features (Yip and Shao, 2021). For example, the HBS1L/PELO complex recognizes the empty ribosomal A site at the 3’ end of a truncated mRNA and promotes ribosome splitting by the recycling factor ABCE1 (Doma and Parker, 2006; Shoemaker et al., 2010). By contrast, attempted translation of poly(A) slows down a leading ribosome to the point that a trailing ribosome collides, resulting in Hel2/ZNF598-mediated ubiquitination of 40S ribosomal proteins and subunit dissociation by the ribosome-associated quality control (RQC)-triggering (RQT) complex (Ikeuchi et al., 2019; Juszkiewicz and Hegde, 2017; Juszkiewicz et al., 2018, 2020; Matsuo et al., 2017, 2020; Simms et al., 2017). An emerging view posits that ribosome collisions comprise a fundamental structural unit recognized by multiple quality control and stress response factors (Kim and Zaher, 2022; Meydan and Guydosh, 2021), including the E3 ligase, Hel2/ZNF598, and the integrated stress response coactivator, GCN1 (Pochopien et al., 2021). Whether there are surveillance mechanisms that specifically respond to stalled ribosomes occupied by translational GTPases, such as eEF1A or release factors, is unknown.

Investigations into how cells recognize and respond to elongation stalls have critically relied on drugs and chemical probes that can modulate translation in a graded, dose-dependent manner (Juszkiewicz et al., 2018, 2020; Simms et al., 2017; Wu et al., 2020; Yan and Zaher, 2021). We recently described ternatin-4, a cyclic peptide that inhibits translation elongation by targeting the complex of eEF1A bound to aminoacyl-tRNAs (aa-tRNA) (Carelli et al., 2015).

Similar to didemnin B (Shao et al., 2016), ternatin-4 stalls elongation by preventing aa-tRNA release from eEF1A on the ribosome (Wang et al., 2022). Based on this mechanism, we reasoned that ternatin-4 could be used as a tool to illuminate cellular responses to elongation-stalled ribosomes in which the GTPase center and A site are occupied by eEF1A bound to aa-tRNA.

Here, we elucidate a novel surveillance pathway, which ultimately results in ubiquitination and degradation of eEF1A trapped on the ribosome by ternatin-4. A CRISPRi screen uncovered two poorly characterized E3 ligases, RNF14 and RNF25, both of which are required for ternatin-induced eEF1A degradation. In response to ternatin-induced stalls, RNF14 and RNF25 play essential roles in the ubiquitination of eEF1A, as well as multiple ribosomal proteins. We further show that GCN1 – whose biological roles outside the canonical ’integrated stress response’ are poorly understood – interacts with RNF14 and is also essential for eEF1A degradation. We propose that this previously undescribed surveillance network – minimally comprising RNF14, RNF25, and GCN1 – monitors the translational status of elongating ribosomes and specifically responds to stalls containing an occluded GTPase center.

## Results

### Ternatin-4 promotes eEF1A degradation in a manner that requires translating ribosomes

In experiments evaluating the cellular response to ternatin-induced elongation stalls, we surprisingly observed a sharp reduction in eEF1A levels after treating HeLa cells with ternatin-4 (**Figure 1A**). Loss of eEF1A was dependent on the concentration of ternatin-4, with an effective half-maximal concentration (EC50) of ∼8 nM. We noted that concentrations higher than 50 nM had progressively diminished effects on eEF1A levels, despite effectively inhibiting translation (**Figure 1B**). This dose-response behavior is reminiscent of mechanistically distinct elongation inhibitors (e.g., cycloheximide, emetine, anisomycin), in which intermediate, but not high concentrations were found to activate ribosome quality control and stress kinase pathways by promoting ribosome collisions (Juszkiewicz et al., 2018; Simms et al., 2017; Wu et al., 2020). Based on these precedents and the observed ‘V-shaped’ dose-response curve (**Figure 1B**), it appeared likely that ternatin-induced ribosome collisions could also play a role in eEF1A degradation.

**Figure 1.**
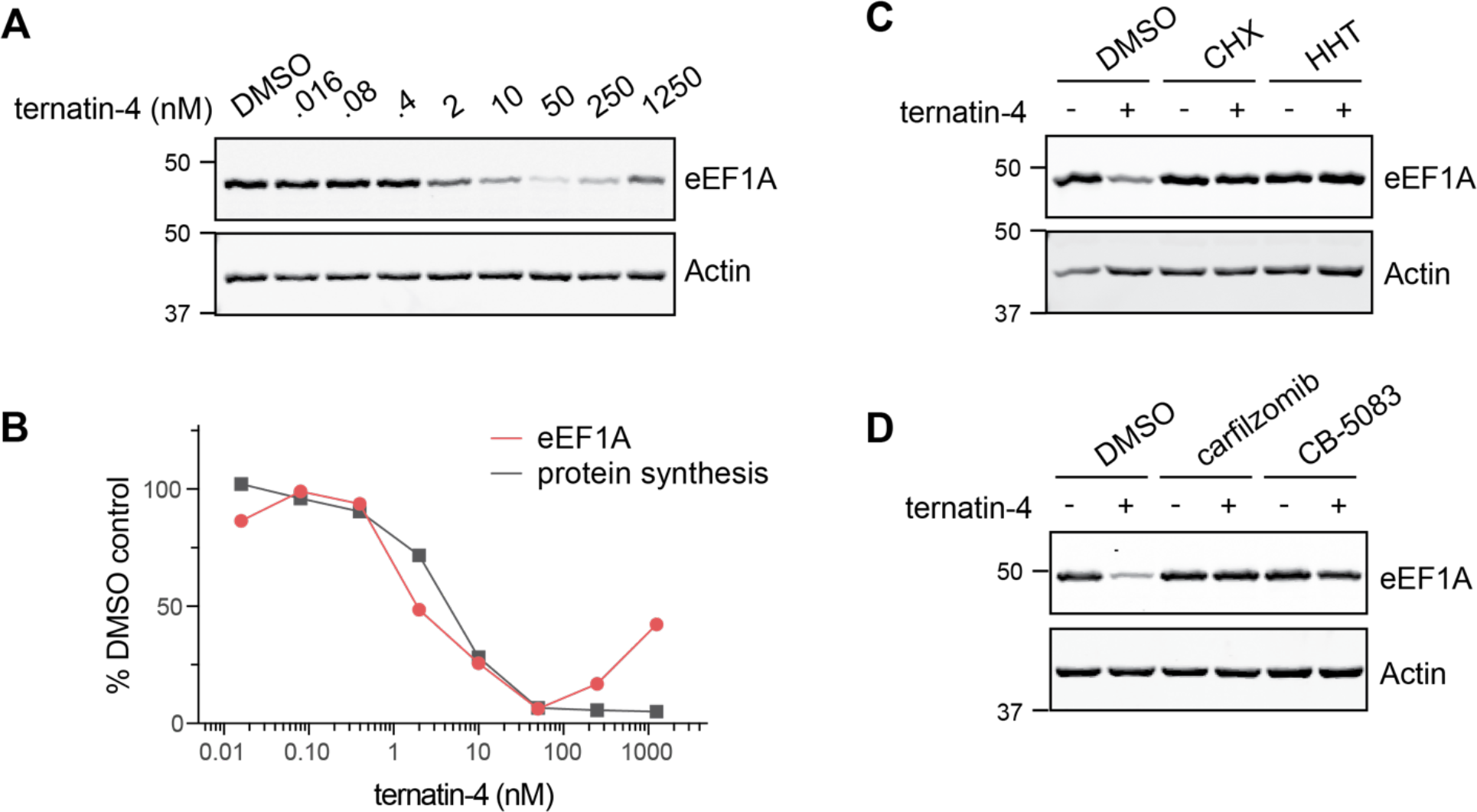
Ternatin-4 promotes eEF1A degradation in a manner that requires translating ribosomes. (**A**) HeLa cells were treated for 20 h with the indicated concentrations of ternatin-4 and analyzed by immunoblotting. (**B**) Quantification of eEF1A levels is derived from (**A**), and protein synthesis measurements were performed on the same day as follows. Following treatment with ternatin-4 for 20 h, HeLa cells were pulsed with O-propargyl puromycin (OPP, 30 μM) for 1 h. Click chemistry with Alexa647-azide was performed on fixed cells to visualize OPP incorporation, and cells were analyzed by flow cytometry. (**C**) HeLa cells were treated for 20 h with ternatin-4 (50 nM) in the presence or absence of cycloheximide (CHX, 50 ug/mL) or homoharringtonine (HHT, 2 μg/mL, 20 min pretreatment). (**D**) HeLa cells were treated for 20 h with ternatin-4 (50 nM) in the presence or absence of the proteasome inhibitor, carfilzomib (500 nM), or the p97 inhibitor, CB-5083 (2.5 μM).

eEF1A is one of the most abundant cellular proteins (∼35 μM in cells) and has been reported to turn over slowly (Zecha et al., 2018). In contrast to results obtained with ternatin-4, treatment of HeLa cells for 20 h with cycloheximide (CHX) or homoharringtonine (HHT) had no discernible effect on eEF1A levels (**Figure 1C**). Hence, inhibiting translation per se is not sufficient to promote loss of eEF1A; rather, direct binding of ternatin-4 to eEF1A on the ribosome might be required. In support of this hypothesis, co-treatment with ternatin-4 and either CHX or HHT – which target the ribosomal E site and peptidyl-transferase center, respectively – completely prevented eEF1A degradation (**Figure 1C**). These results suggest that ternatin-induced eEF1A degradation requires actively translating ribosomes that are competent to bind eEF1A/aa-tRNA; such ribosomes would be less abundant in cells treated with cycloheximide or homoharringtonine owing to their inhibitory effects on mRNA-tRNA translocation and initiation, respectively (Budkevich et al., 2011; Fresno et al., 1977). Ternatin-induced eEF1A degradation was also prevented by co-treatment with either proteasome or p97/VCP inhibitors (**Figure 1D**), thus implicating the ubiquitin/proteasome system (UPS) in this process. Collectively, our findings suggest that ternatin-4, which traps eEF1A on the ribosome and prevents aa-tRNA accommodation into the A site (Wang et al., 2022), targets eEF1A for UPS-mediated destruction by a mechanism that requires elongation-competent ribosomes.

### CRISPRi screen reveals two E3 ligases required for ternatin-induced eEF1A degradation

We sought to identify the UPS components, and in particular the E3 ligase(s) required for ternatin-induced eEF1A degradation. To do this, we performed a CRISPRi screen using an mCherry-eEF1A fusion as a fluorescent reporter, which also contains an internal ribosome entry site followed by GFP. We stably expressed this bicistronic reporter construct, along with dCas9- BFP-KRAB, in HCT116 cells that are homozygous for an A399V mutation in the didemnin/ternatin-4 binding site of eEF1A1. We previously showed that this mutation abrogates ternatin-4 binding and confers complete resistance to its cellular effects (Carelli et al., 2015).

Hence, this reporter cell line, which expresses mCherry-eEF1A at low levels relative to endogenous A399V eEF1A, allows us to study ternatin-induced eEF1A degradation without globally inhibiting protein synthesis. Treatment of the reporter cells with ternatin-4 resulted in a strong reduction in mCherry-eEF1A fluorescence, whereas GFP was unaffected (**Figure 2A**). Loss of mCherry-eEF1A was dependent on the ternatin-4 concentration (EC50 ∼5 nM; **Figure S1A**) and treatment time (t1/2 ∼5 h; **Figure S1B**), and it was prevented by co-treatment with either a proteasome inhibitor or ribosome-targeted translation inhibitors (**Figure S1C**), similar to endogenous eEF1A (**Figure 1**). Importantly, ternatin-4 had no effect on mCherry-eEF1A bearing the A399V mutation, which confirms that ternatin-4 binding is required to promote eEF1A degradation (**Figure S1D**).

**Figure 2.**
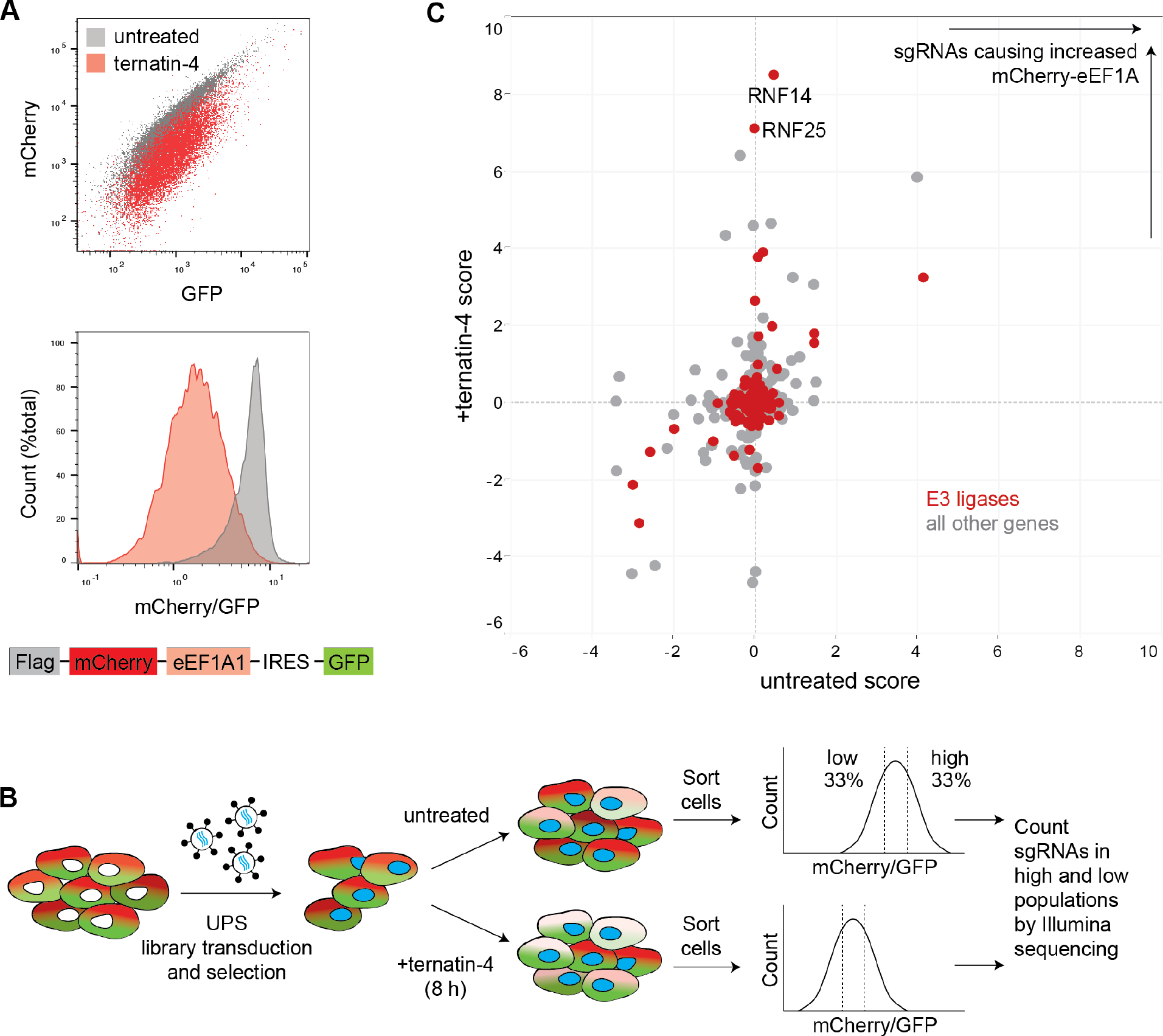
CRISPRi screen reveals two E3 ligases required for ternatin-induced eEF1A degradation . (**A**) Fluorescent reporter to monitor eEF1A degradation. Ternatin-resistant HCT116 cells (eEF1A1^A399V/A339V^) stably expressing mCherry-eEF1A1_IRES-GFP and dCas9-HA-NLS-KRAB- BFP were treated ± ternatin-4 (50 nM) for 8 h and analyzed by flow cytometry. (**B**) Schematic of CRISPRi screen. Cells from (**A**) were transduced with a library targeting ∼1700 genes (5 sgRNAs/gene) related to ubiquitin signaling and proteostasis. After 2 days, cells were selected with puromycin for 72 h, allowed to recover for 48 h, and treated with 50 nM ternatin-4 for 8 h or left untreated. Cells were sorted into high and low mCherry/GFP populations, and sgRNA counts were determined by deep sequencing. (**C**) CRISPRi scores (based on sgRNA enrichment in high vs. low mCherry/GFP populations) from cells treated with ternatin-4 (Y-axis) or left untreated (X-axis). Knockdown of *RNF14* or *RNF25* stabilizes mCherry-eEF1A1 levels in ternatin-treated but not untreated cells. Scores (plotted for each gene) are the product of the phenotype value (log2 of the average high/low sgRNA counts for the three most extreme sgRNAs per gene) and the negative log10 of the p value.

We transduced the mCherry-eEF1A reporter cells with a focused sgRNA library targeting ∼1700 genes primarily involved in ubiquitin signaling and proteostasis (5 sgRNAs/gene, see **Table S1** for the complete list of genes), plus 250 nontargeting sgRNAs (Chen et al., 2019). Transduced cells were treated with ternatin-4 for 8 h or left untreated, followed by sorting into high and low fluorescence populations based on mCherry normalized to GFP (**Figure 2B**). For each population, sgRNA frequencies were quantified by Illumina sequencing and analyzed using our established bioinformatics pipeline (Kampmann et al., 2013; Tian et al., 2019). We focused on genes whose knockdown led to increased mCherry-eEF1A levels (less degradation) in ternatin-treated but not untreated cells. Two E3 ligase genes, *RNF14* and *RNF25*, emerged as the most prominent hits (**Figure 2C**).

Knockdown of either *RNF14* or *RNF25* strongly prevented eEF1A degradation, with multiple sgRNAs enriched in the high mCherry-eEF1A population of ternatin-treated (but not untreated) cells (**Figure S1E**). By contrast, sgRNAs targeting previously characterized ribosome-associated E3 ligases and RQC factors – including *ZNF598*, *LTN1*, *RNF10*, and *CNOT4* – had little or no effect on eEF1A degradation (**Figure S1F** and **Table S1**). To validate *RNF14* and *RNF25*, we retested individual, top-scoring sgRNAs in our mCherry-eEF1A reporter cells and observed a complete loss of ternatin-induced degradation (**Figure 3A**).

**Figure 3.**
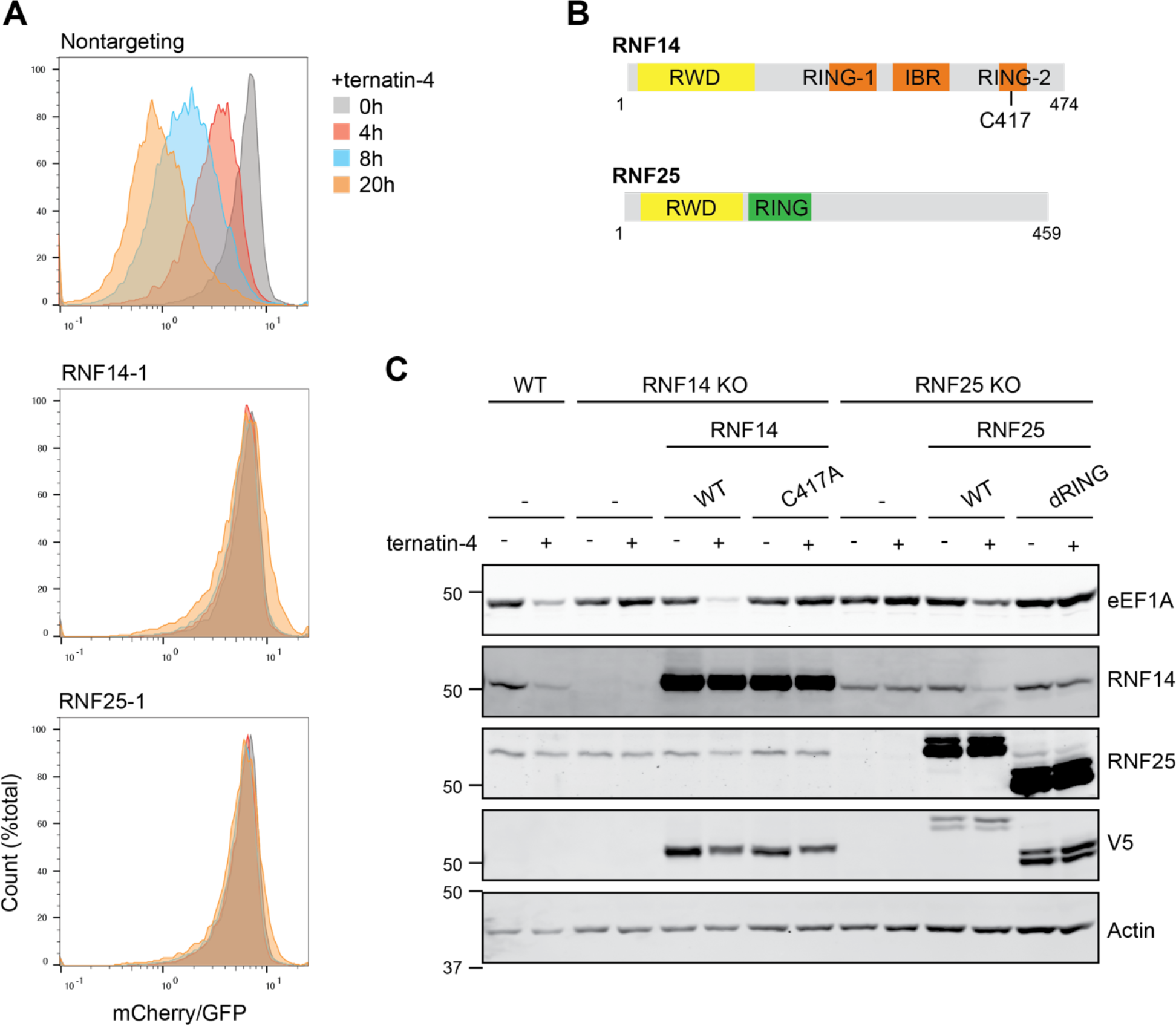
RNF14 and RNF25 are required for ternatin-induced eEF1A degradation. (**A**) CRISPRi screening cells were transduced with the top-scoring *RNF14 and RNF25* sgRNAs from the screen in Figure 2. Transduced cells were treated with 50 nM ternatin-4 for the indicated times and analyzed by flow cytometry. (**B**) Domain organization of RNF14 and RNF25. The RNF14 catalytic cysteine (C417) in the RING-2 domain is indicated. (**C**) RNF14 and RNF25 HeLa KO cells were generated using CRISPR/Cas9. Cells were further transduced with the indicated V5-tagged RNF14 or RNF25 constructs (both with IRES-mCherry) and sorted to produce pure populations. Cells were treated for 20 h with ternatin-4 (50 nM) and analyzed by immunoblotting.

RNF14 (also called ARA54) (Kang et al., 1999) belongs to the Ring-Between-Ring (RBR) class of E3 ligases, which transfer ubiquitin from an E2 (bound to the RING1 domain) to a conserved catalytic cysteine (Cys417 in RNF14) in the RING2 domain (Dove and Klevit, 2017; Walden and Rittinger, 2018). By contrast, RNF25 (also called AO7) (Lorick et al., 1999) is a RING-type E3 ligase. Deletion of RNF25 was recently found to sensitize cells to the alkylating agent, methyl methanesulfonate (MMS) (but not other DNA damaging agents), indicating a potential role in repairing methylated DNA (Hundley et al., 2021). We note that MMS also methylates RNA and was found to activate the RQC pathway in yeast (Yan and Zaher, 2021), suggesting that MMS hypersensitivity of RNF25 KO cells could also stem from effects related to translation. Both RNF14 and RNF25 contain N-terminal RWD domains (**Figure 3B**), protein interaction domains found in ∼30 diverse human proteins exemplified by the ribosome-associated stress kinase, GCN2. Overall, the biological functions of RNF14 and RNF25 remain poorly understood.

To further validate RNF14 and RNF25, we generated HeLa knockout cell lines using CRISPR/Cas9. Similar to our CRISPRi knockdown results with mCherry-eEF1A, knockout of either RNF14 or RNF25 abolished ternatin-induced degradation of endogenous eEF1A, while having no effect on eEF1A levels in untreated cells (**Figure 3C**). This phenotype was consistently observed across multiple RNF14 and RNF25 KO clones (**Figure S2A**).

Concomitant with eEF1A degradation, ternatin-4 treatment of wild type HeLa cells caused a dramatic reduction in endogenous RNF14 levels (**Figure 3C** and **Figure S2B**). Remarkably, this effect was also abolished in RNF25 KO cells (**Figure 3C**). These results suggest: (1) RNF14, while an essential mediator of eEF1A destruction, is also degraded in response to ternatin-4 treatment, and (2) RNF25 plays an essential role in ternatin-induced degradation of both eEF1A and RNF14. While the precise biochemical choreography remains to be elucidated, we hypothesize that RNF14 and RNF25 work together as part of a ubiquitin-dependent signaling network that senses and responds to ternatin-induced elongation stalls (see Discussion).

Reintroduction of wild type RNF14 and RNF25 into their respective KO cells restored eEF1A degradation, confirming an essential role for both E3 ligases (**Figure 3C**).

Overexpression of wild type RNF14 rescued and further enhanced eEF1A degradation in the KO cells, whereas overexpression of the catalytic Cys417 to Ala mutant failed to restore degradation. Likewise, expression of wild type RNF25, but not a RING deletion mutant, restored degradation of both eEF1A and RNF14 in the RNF25 KO cells (**Figure 3C**). Overexpression of RNF14, but not RNF25, in wild type cells also led to enhanced eEF1A degradation, whereas a catalytically dead mutant of RNF14 (but not RNF25) acted in a dominant negative manner, completely abrogating eEF1A degradation (**Figure S2C**). These results suggest that the ubiquitin ligase activity of RNF14 is rate-limiting for eEF1A degradation and that overexpression of catalytically dead RNF14 can saturate a binding site required to promote eEF1A degradation.

### RNF14 and RNF25 mediate ubiquitination of eEF1A and ribosomal proteins

Having established the essential roles of RNF14 and RNF25 E3 ligase activity in eEF1A degradation, we sought to determine the global landscape of RNF14/RNF25-dependent ubiquitination sites in an unbiased manner. To do this, we used an established SILAC proteomics workflow in which tryptic peptides containing a ubiquitin-derived diGly remnant (attached to a substrate lysine residue) are immuno-enriched and quantified by mass spectrometry (Kim et al., 2011). We first treated cells stably overexpressing RNF14 with or without ternatin-4 for 4 h in two biological replicates (SILAC light/heavy label swaps; see Methods). Cell lysates from each treatment condition (± ternatin-4) were combined and digested with trypsin, prior to enrichment of diGly peptides (**Figure S3A**). Mass spectrometry analysis revealed >800 ubiquitination sites, 7 of which were strongly and reproducibly induced by ternatin (log2-fold change >2) in both replicates (**Figure 4A**). Strikingly, all 7 top-ranked sites were found either in eEF1A (4 sites) or the ribosomal proteins RPLP0, RPS13, and RPS17.

**Figure 4.**
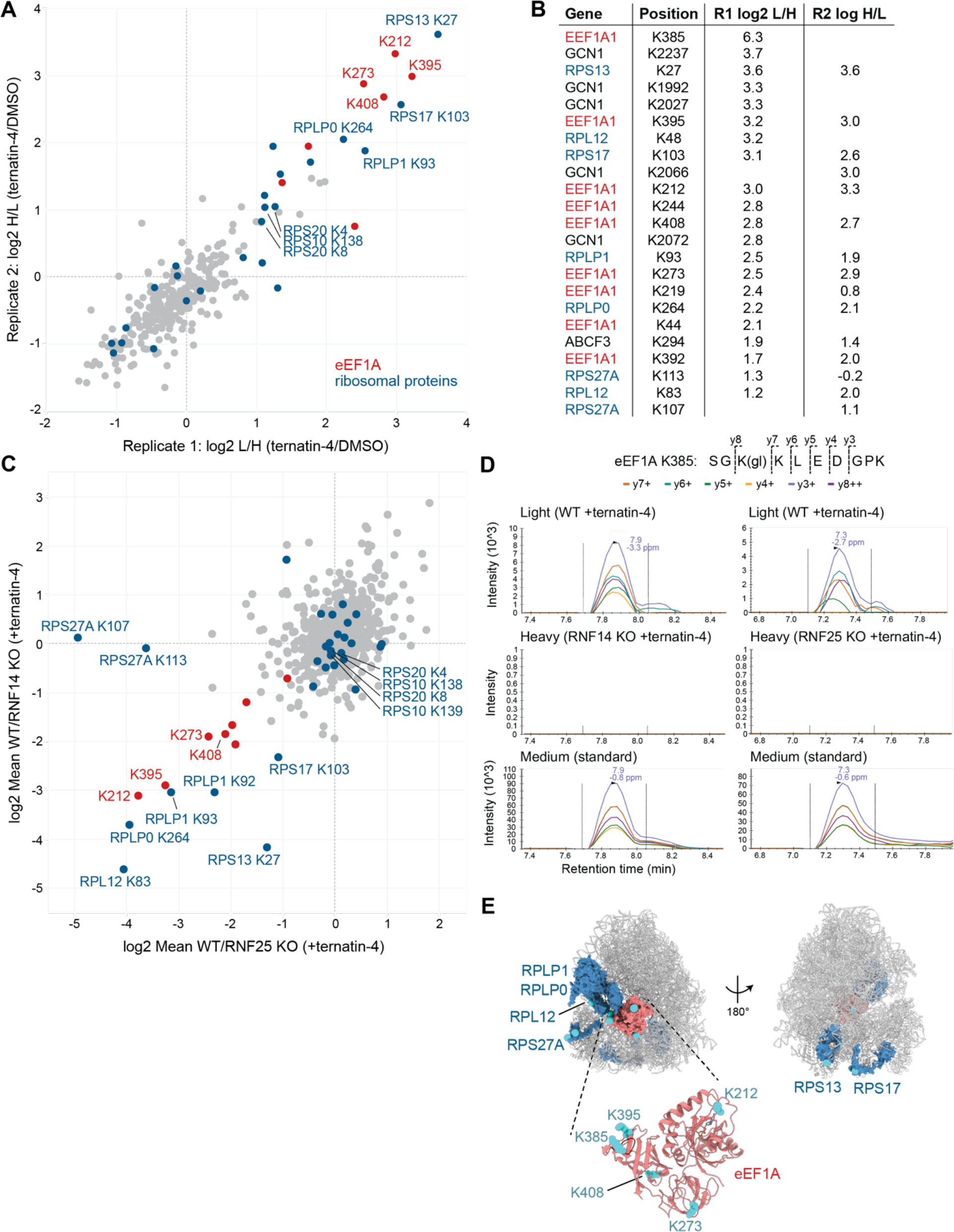
Ternatin-4 promotes RNF14 and RNF25-dependent ubiquitination of eEF1A and ribosomal proteins. (**A**) HeLa cells stably overexpressing RNF14 were labeled in SILAC media (containing heavy or light Arg/Lys) and treated for 4 h with 50 nM ternatin-4 or DMSO. Lysates from ternatin-4 and DMSO-treated cells were mixed, trypsinized, and desalted. Peptides containing diGly ubiquitin remnants were immuno-enriched and analyzed by LC-MS/MS. Shown are SILAC ratios for each diGly site identified in both biological replicates. (**B**) SILAC ratios for top-ranked diGly sites (eEF1A and ribosome-associated proteins) from each of the two biological replicates from part (**A**). (**C**) HeLa WT and RNF14 or RNF25 KO cells were treated for 4 h with 50 nM ternatin-4. KO and WT SILAC pairs were mixed before processing for LC-MS/MS analysis as described in (**A**). Mean KO/WT SILAC ratios were calculated from two biological replicates (H/L label swaps) for both RNF14 KO and RNF25 KO experiments. (**D**) Samples from (**C**) were analyzed by PRM-MS for eEF1A K385 ubiquitination. Chromatograms are shown for one biological replicate each for RNF14 and RNF25 KO samples. MS2 transitions were monitored for light (WT cells), heavy (KO cells), and medium (synthetic peptide standard) variants of SGK[diGly]KLEDGPK, corresponding to eEF1A K385-Ub. (**E**) Cartoon showing location of ternatin-induced and RNF14/25-dependent ubiquitination sites (cyan spheres) on eEF1A and ribosomal proteins based on cryo-EM models (PDB: 5LZS; RPLP0/RPLP1 model based on PDB: 4V6X). The loop containing eEF1A K385 (residues 379-386) is colored in dark red in the ribbon model.

Several ubiquitination sites in proteins relevant to translation elongation – including 7 additional eEF1A sites – were induced to a somewhat lower extent (log2-fold change >1) or were identified in only one biological replicate (**Figure 4B** and **Table S2**), as is typical in data-dependent acquisition (DDA) mass spectrometry. Of all identified ubiquitination sites, eEF1A K385 was induced most strongly by ternatin, increasing by at least 75-fold (**Figure 4B** and **S3B**). In addition, ternatin-4 treatment led to increased ubiquitination of multiple sites within the C-terminal region of the ribosome collision sensor, GCN1 (**Figure 4B** and **S3B**), as well as ABCF3 (ortholog of yeast Gcn20) and the ribosomal proteins, RPL12 and RPS27A (**Figure 4B**). We repeated the diGly SILAC-MS experiment (± ternatin-4) in RNF14-overexpressing cells co-treated with a proteasome inhibitor and in wild type cells (without proteasome inhibitor treatment), and we observed similar levels of ternatin-induced eEF1A and ribosomal protein ubiquitination (**Figures S3B** and **S3C, Table S2**).

We next employed SILAC-MS to quantify ternatin-induced ubiquitination sites in RNF14 or RNF25 KO cells, relative to the parental wild type cells. Consistent with their obligate roles in eEF1A degradation, knockout of either RNF14 or RNF25 dramatically reduced eEF1A ubiquitination at multiple sites (**Figure 4C** and **Table S2**). Ubiquitination of a discrete set of ribosomal protein sites, including RPLP0 K264, RPLP1 K92, RPLP1 K93, and RPL12 K83, was similarly reduced in both knockout cell lines. In striking contrast, ubiquitination of RPS27A at K107 and K113 was reduced in RNF25 KO but not RNF14 KO cells, suggesting that these ubiquitination events may be selectively mediated by RNF25 (**Figure 4C**). Ubiquitination of ZNF598-dependent sites on RPS10 and RPS20 (Garzia et al., 2017; Juszkiewicz et al., 2018; Sundaramoorthy et al., 2017) was unaffected by RNF14 or RNF25 knockout (**Figure 4C** and **Table S2**), despite increasing ∼2-fold in response to ternatin-4 treatment (**Figure 4B** and **Table S2**). These data suggest that ternatin-induced stalls can activate distinct ribosome-associated ubiquitination events, which are mediated independently by ZNF598 and RNF14/RNF25.

The ubiquitination site most strongly induced by ternatin-4, eEF1A K385, was not identified in every biological replicate; this is likely due to the presence of a missed trypsin cleavage site after K386 (the fully trypsinized tetrapeptide would be too short to identify unambiguously). However, across multiple independent SILAC-MS experiments in which it was identified, K385 ubiquitination was consistently and dramatically induced by ternatin treatment, suggesting its likely dependence on RNF14/RNF25. To test this rigorously, we spiked in a synthetic ’medium heavy’ peptide standard (SGK[diGly]K<u>L</u>EDGPK, synthesized with heavy Leu), which facilitated identification and quantification of the endogenous SILAC-derived peptides (SG<u>K</u>[diGly]<u>K</u>LEDGP<u>K</u>, containing 3 light or heavy Lys) via parallel reaction monitoring (PRM) mass spectrometry instead of DDA-MS (see Methods for details). These experiments unambiguously revealed diGly-modified eEF1A K385 in ternatin-treated wild type cells, whereas it was undetectable in either RNF14 or RNF25 knockout cells (**Figure 4D**).

We conclude that ternatin-induced elongation stalls promote ubiquitination of multiple sites on eEF1A and a discrete set of ribosomal proteins. Most ternatin-induced ubiquitination sites are dependent on both RNF14 and RNF25, whereas RPS27A ubiquitination selectively requires RNF25. Several of the ribosomal ubiquitination sites we identified – including those on RPLP0, RPLP1, RPL12, and RPS27A – are proximal to the GTPase center where eEF1A binds, consistent with the notion that RNF14/RNF25-dependent eEF1A ubiquitination occurs on elongation-stalled ribosomes (**Figure 4E**). By contrast, RPS13 and RPS17 sites – which are also ternatin induced and RNF14/25 dependent – localize to a distinct region of the 40S subunit near the interface of collided di-ribosomes. Whether ribosomal protein ubiquitination contributes in some way to eEF1A degradation or plays some other role remains to be determined.

### K385 is required for efficient eEF1A degradation and is directly ubiquitinated by RNF14

To assess the functional relevance of ternatin-induced, RNF14/RNF25-dependent eEF1A ubiquitination sites identified by mass spectrometry, we introduced lysine to arginine mutations into our mCherry-eEF1A reporter. With the exception of K385, mutation of individual lysines had little or no effect on eEF1A degradation kinetics. By contrast, ternatin-induced degradation of K385R eEF1A was impaired (**Figure 5A** and **5B**). K385 resides on a long, flexible loop in the C-terminal beta-barrel domain of eEF1A (**Figure 4E**). This loop lies near the interface between the C-terminal domain and the N-terminal GTPase domain of ribosome-bound eEF1A and directly contacts the ternatin/didemnin binding site (Carelli et al., 2015; Shao et al., 2016). The mutagenesis results suggest that K385 ubiquitination, which is dramatically increased in the context of ternatin-induced stalls, plays a critical role in subsequent events leading to p97/VCP and proteasome-dependent eEF1A degradation. Additional ternatin-induced ubiquitination sites, as revealed by our SILAC-MS experiments, may also contribute collectively to promote efficient eEF1A degradation.

**Figure 5.**
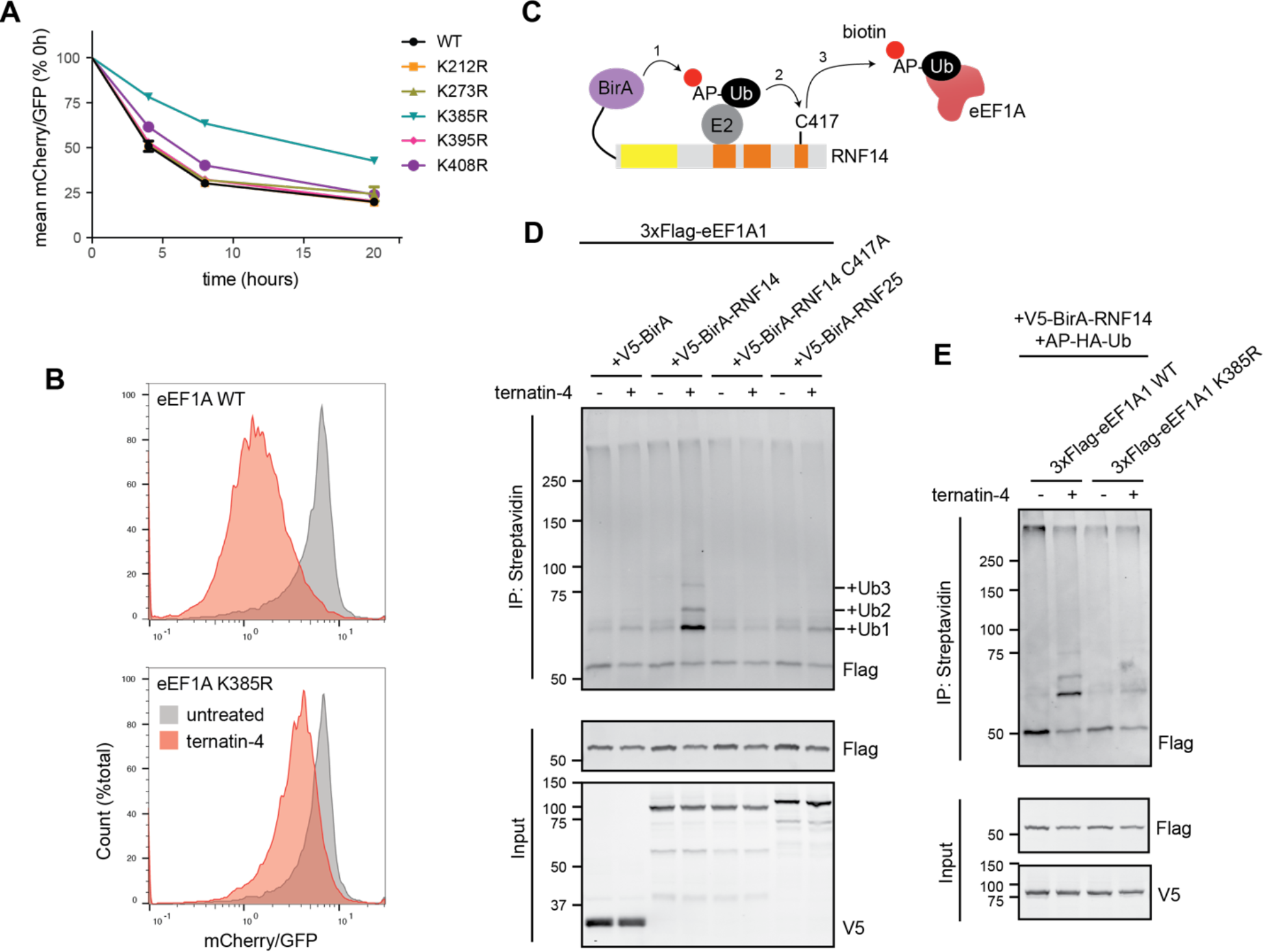
eEF1A K385 is required for efficient degradation and is directly ubiquitinated by RNF14. (**A**) Cells expressing WT and mutant mCherry-eEF1A reporter constructs were treated with ternatin-4 (50 nM) for the indicated times and analyzed by flow cytometry. Plotted are the mean mCherry/GFP ratios (n=3, ± SEM) relative to t=0 h for each mutant. (**B**) Histograms for cells from (**A**) expressing WT or K385R eEF1A and treated ± ternatin-4 for 8 h. (**C**) Schematic for proximity-based biotin transfer from E3 (BirA-RNF14) to ubiquitin/E2 to substrates (eEF1A). Acceptor peptide (AP)-ubiquitin is biotinylated when bound to E2 and the biotin ligase BirA- RNF14 fusion. Biotinylated AP-Ub is subsequently transferred to E3 substrates, which are enriched via streptavidin. (**D**) Ternatin-4 induces proximity-mediated eEF1A ubiquitination by RNF14, but not RNF25. Ternatin-resistant HCT116 cells stably expressing 3xFlag-eEF1A1 were co-transfected with AP-HA-Ub and the indicated BirA fusion constructs. Cells were treated with 50 μM biotin ± ternatin-4 (50 nM) for 4 h. (**E**) As in part (**D**), except cells expressing WT or K385R 3xFlag-eEF1A were transfected with AP-HA-Ub and BirA-RNF14 and treated with biotin ± ternatin-4.

Based on the genetic evidence implicating RNF14 and RNF25 in ternatin-induced eEF1A ubiquitination and degradation, we hypothesized that one or both E3 ligases directly ubiquitinate eEF1A K385. To test whether this occurs in cells, we turned to a biotin-transfer assay in which an E3 of interest is fused to the biotin ligase BirA (Deshar et al., 2016, 2019; Yoo et al., 2019). This proximity-based biotinylation assay relies on the specificity of BirA for an acceptor peptide fused to ubiquitin (AP-Ub). Expression of the BirA-E3 fusion protein results in biotinylation of a proximal E2-bound AP-Ub, followed by transfer of biotin-AP-Ub to a proximal substrate (**Figure 5C**). Enrichment of biotinylated proteins with streptavidin-conjugated beads allows detection of ubiquitinated substrates specific to the BirA-E3 ligase of interest.

We established the BirA-E3 proximity biotinylation assay in our ternatin-resistant HCT116 cell line (eEF1A1^A399V/A339V^), modified to stably express Flag-eEF1A. Strikingly, treatment of cells co-expressing BirA-RNF14 and AP-Ub with ternatin-4 for 4 h resulted in the specific enrichment of one major (mono-Ub) and two minor (di- and tri-Ub) higher-molecular weight forms of Flag-eEF1A (**Figure 5D** and **S4A**). Transfer of biotin-AP-Ub to Flag-eEF1A was not observed in cells expressing similar levels of BirA or an active site mutant of BirA-RNF14 (C417A), and it was barely detectable in cells expressing BirA-RNF25 (**Figure 5D**). Finally, we observed a drastic decrease in BirA-RNF14-mediated ubiquitination of K385R Flag-eEF1A (**Figure 5E** and **S4B**). These results suggest that RNF14 can directly promote eEF1A K385 ubiquitination in response to ternatin-induced elongation stalls. By contrast, the essential role of RNF25 in promoting eEF1A K385 ubiquitination, as revealed by our PRM assay (**Figure 4C**), is likely indirect and may be upstream of RNF14 (see Discussion).

### GCN1 interacts with RNF14 and is essential for ternatin-induced eEF1A degradation

Besides eEF1A and ribosomal proteins, our diGly proteomics experiments revealed multiple ternatin-induced ubiquitination sites on GCN1. GCN1 is a conserved ribosome-associated scaffolding protein that binds the RWD domain of the integrated stress response kinase, GCN2 (Sattlegger and Hinnebusch, 2000; Kubota et al., 2000). In addition, GCN1 was recently shown to interact with collided, elongation-stalled ribosomes in yeast (Pochopien et al., 2021) and human cells (Wu et al., 2020). Given its established interactions with stalled ribosomes, we postulated that GCN1 (which was not included in the CRISPRi screen) might play a direct role in RNF14/RNF25-mediated eEF1A degradation.

To evaluate potential interactions with GCN1, we immunoprecipitated Flag-tagged RNF14 or RNF25 (stably expressed in HeLa cells). Because previous studies employed chemical crosslinking to stabilize protein interactions with GCN1 (Sattlegger and Hinnebusch, 2005; Wu et al., 2020), we briefly treated cells with 0.1% formaldehyde (22 °C, 10 min) prior to preparing detergent lysates for immunoprecipitation. This experiment revealed a specific interaction between Flag-RNF14 and endogenous GCN1 in cells treated with or without ternatin (**Figure 6A**). A higher-molecular weight, likely multi-ubiquitinated form of GCN1 was specifically enriched in Flag-RNF14 immunoprecipitates from ternatin-treated cells, whereas this higher-molecular weight GCN1 species was not detected in immunoprecipitates from untreated cells or from ternatin-treated cells expressing a catalytically dead mutant (C417A) of Flag-RNF14 (**Figure 6A** and **S5A**). Relative to Flag-RNF14, Flag-RNF25 enriched GCN1 to a lesser extent, yet robustly immunoprecipitated endogenous 40S and 60S ribosomal proteins, even in the absence of chemical crosslinking (**Figure 6A**). These results suggest that RNF14 and RNF25 can form complexes containing GCN1 and ribosomes, consistent with a requirement for both E3 ligases in ternatin-induced ubiquitination of eEF1A and ribosomal proteins (**Figure 4**) and the established role of GCN1 as a sensor for elongation-stalled ribosomes (Meydan and Guydosh, 2020; Pochopien et al., 2021; Wu et al., 2020; Yan and Zaher, 2021). The data also suggest that RNF14 may directly ubiquitinate GCN1, although the functional relevance of these marks remains to be determined.

**Figure 6.**
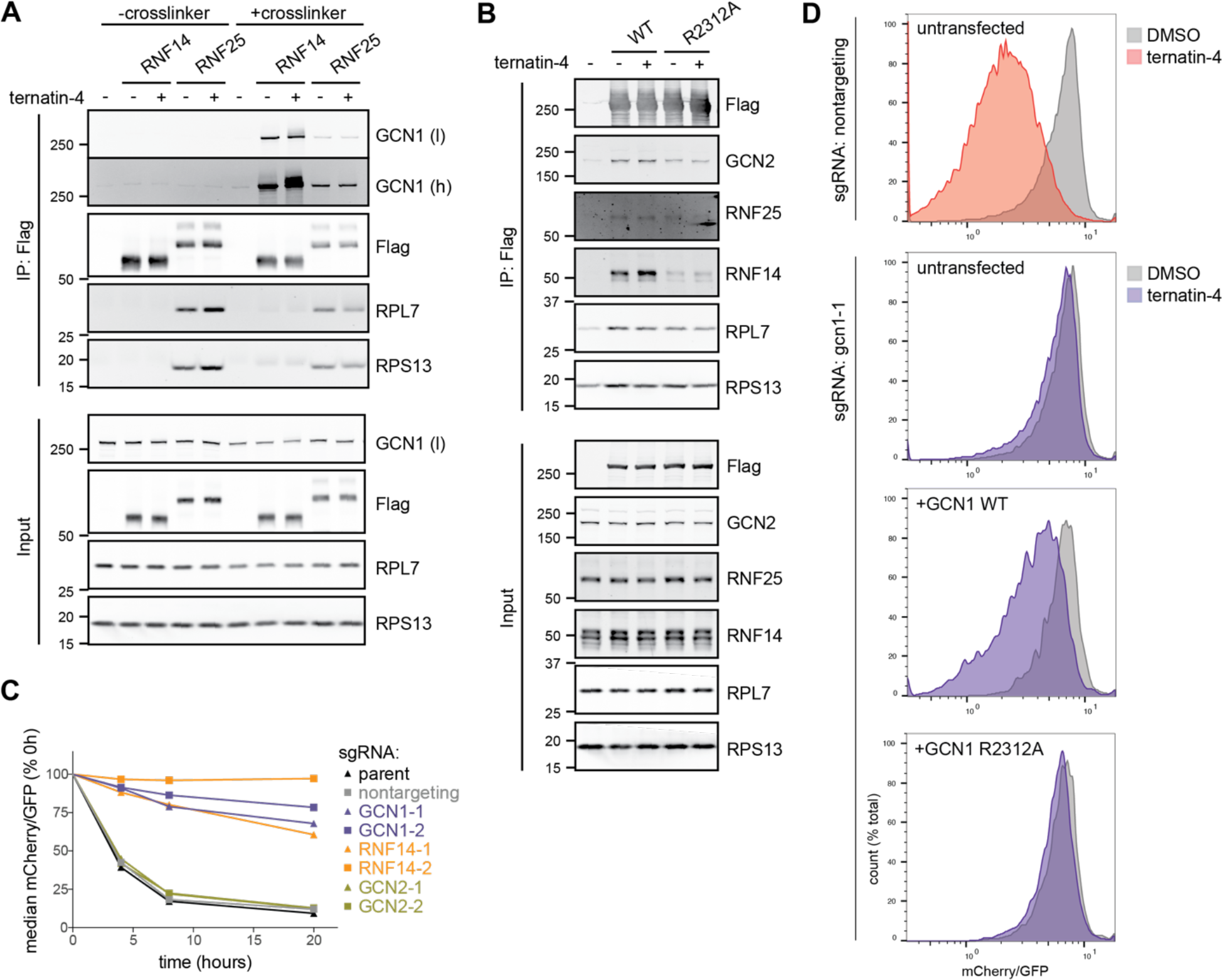
GCN1 interacts with RNF14 and is essential for ternatin-induced eEF1A degradation. (**A**) HeLa cells stably expressing 3xFlag-tagged RNF14 or RNF25 were treated for 4 h ± ternatin-4 (50 nM) and then crosslinked with 0.1% PFA (10 min, 22 °C) prior to cell lysis, Flag immunoprecipitation, and immunoblotting. (**B**) HEK293T cells transiently transfected with WT or R2312A GCN1-3xFlag were treated and analyzed as in part (**A**). (**C**) mCherry-eEF1A/CRISPRi reporter cells were transduced with sgRNAs targeting the indicated genes as in Figure 3. Cells were treated with ternatin-4 (50 nM) for the indicated times, and mCherry-eEF1A1 levels were analyzed by flow cytometry. (**D**) mCherry-eEF1A/CRISPRi reporter cells were first transduced with nontargeting or GCN1-targeted sgRNAs as in (**C**). Cells were then transfected with WT or R2312A GCN1-3xFlag (or untransfected), treated for 8 h ± ternatin-4 (50 nM), and analyzed by flow cytometry.

GCN1-interacting proteins other than GCN2 remain relatively unexplored in higher eukaryotes. By contrast, a conserved C-terminal region of yeast Gcn1, which includes R2259, has been shown to mediate interactions with the RWD domains of yeast Gcn2, Yih1, and Gir2 (Castilho et al., 2014; Pochopien et al., 2021). Mutation of Gcn1 R2259 (corresponding to R2312 in human GCN1) had no effect on binding to ribosomes or Gcn20, yet abolished interactions with Gcn2 and prevented Gcn2 activation upon amino acid starvation (Sattlegger and Hinnebusch, 2000). Like GCN2, both RNF14 and RNF25 contain RWD domains, which we hypothesized could mediate interactions with the conserved GCN2-binding region of GCN1.

Consistent with the above results using overexpressed RNF14/RNF25 (**Figure 6A**), reciprocal immunoprecipitation of overexpressed GCN1-Flag confirmed binding to endogenous RNF14, whereas binding to endogenous RNF25 was not reliably detected over background (**Figure 6B**). Importantly, mutation of R2312 in the conserved RWD binding region of GCN1 dramatically reduced interaction with RNF14, but not ribosomal proteins. GCN2 binding was likewise diminished with GCN1 R2312A, although to a lesser extent than RNF14.

To test for a functional role of GCN1 in eEF1A degradation, we used CRISPRi and the mCherry-eEF1A reporter assay. Remarkably, GCN1 knockdown diminished ternatin-induced eEF1A degradation to a similar extent as RNF14 knockdown (**Figures 6C** and **S5B**), whereas knockdown of the canonical GCN1 partner, GCN2, had no effect. Impaired eEF1A degradation was partially rescued upon transfection of the CRISPRi knockdown cells with wild type GCN1. By contrast, the RNF14 binding-defective GCN1 mutant (R2312A) completely failed to restore eEF1A degradation in cells depleted of endogenous GCN1 (**Figure 6D** and **S5C**). Collectively, the results in **Figure 6** demonstrate that GCN1, which has previously been linked to activation of the integrated stress response kinase GCN2, forms a distinct functionally relevant complex with the E3 ligase RNF14. Based on the strong inhibitory effect of the R2312A mutation, this interaction is likely mediated by the N-terminal RWD domain of RNF14, in direct competition with other GCN1-binding proteins such as GCN2 and IMPACT. We propose that RNF14 (and possibly RNF25; see Discussion) is recruited by GCN1 to stalled ribosomes and locally activated, resulting in eEF1A ubiquitination and ultimately, proteasome and p97/VCP-dependent degradation (**Figure 7A**). To our knowledge, this is the first demonstration that GCN1, a sensor for elongation-stalled ribosomes, plays an essential role in an E3 ligase network.

**Figure 7.**
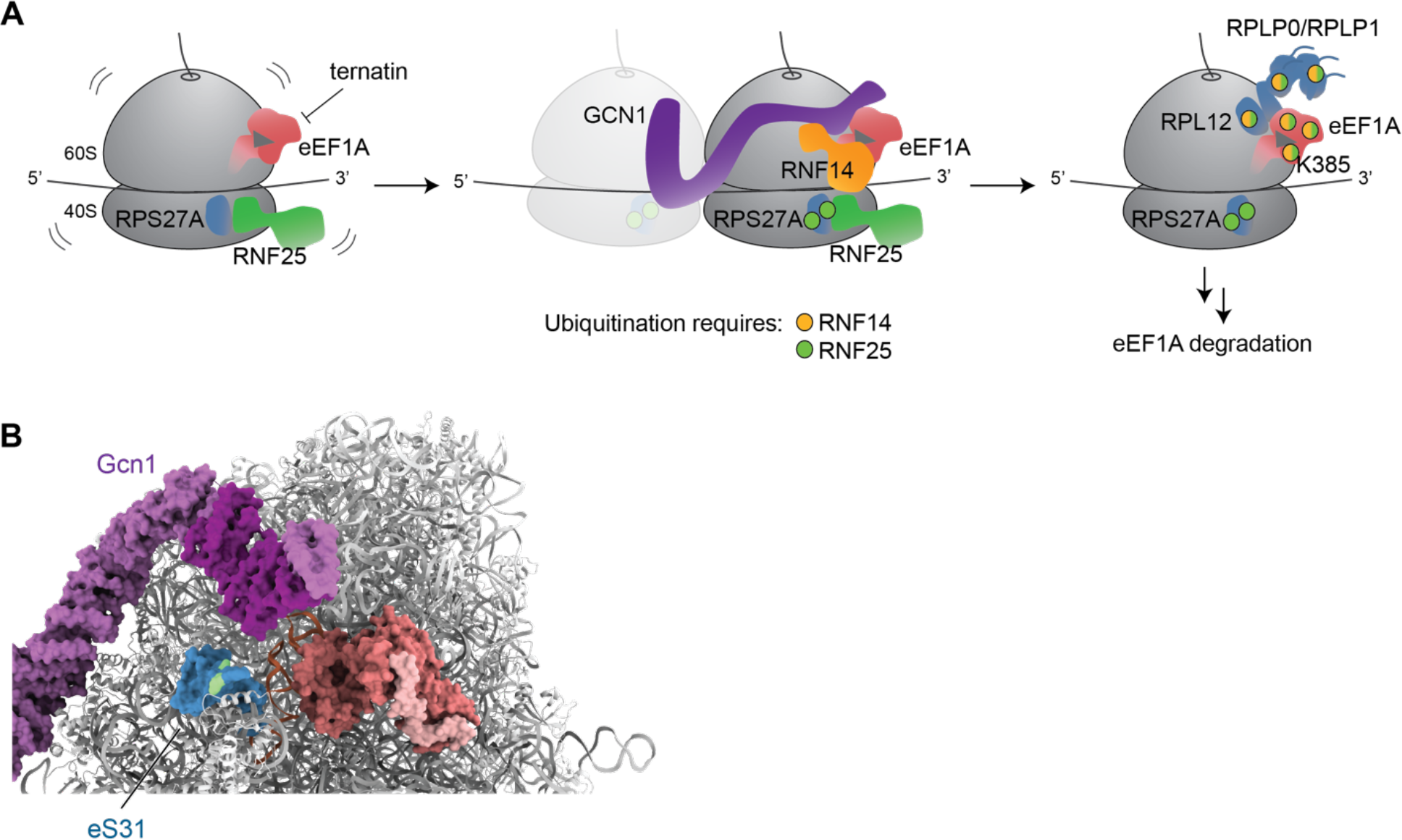
Model for elongation stall-induced activation of the GCN1/RNF14/RNF25 pathway. (**A**) RNF25 binds ribosomes in unstressed cells and ubiquitinates the 40S ribosomal protein RPS27A, possibly in response to infrequent and/or transient elongation pausing events. Upon ternatin-induced stalling, GCN1-bound RNF14 is recruited to ribosome collisions, leading to ubiquitination of trapped eEF1A and the adjacent ribosomal proteins RPL12, RPLP0, and RPLP1. Ubiquitination of eEF1A at K385 and other sites leads to p97 and proteasome- dependent eEF1A degradation. (**B**) CryoEM structure of yeast Gcn1/di-ribosome complex (PDB: 7NRC) showing Gcn1 (purple) bound to the leading (stalled) ribosome. The conserved RWD- binding region of Gcn1 (magenta) is proximal to both eS31 (blue; ortholog of human RPS27A/eS31) and the GTPase center, which in this structure contains the yeast Gir2/Rbg2 complex (pink/red) bound to peptidyl-tRNA in the A site. Maximal RNF14 E3 ligase activity may involve allosteric interactions with GCN1 (magenta region) and/or RNF25-dependent ubiquitin marks on RPS27A K107/K113 (green), by analogy to other RBR-type E3s (e.g., parkin and HOIP).

## Discussion

We investigated whether elongation stalls induced by ternatin-4, which traps eEF1A on the ribosome and prevents aa-tRNA accommodation (Wang et al., 2022), would activate as yet uncharacterized quality control pathways. We discovered that ternatin-4 robustly induces eEF1A degradation, and we used this phenotype as a foothold to elucidate a novel QC/stress response pathway requiring two poorly characterized E3 ligases, RNF14 and RNF25. RNF14 and RNF25 act in concert with the ribosome collision sensor GCN1 – best known as an upstream activator of the integrated stress response kinase GCN2 – to promote ubiquitination of eEF1A and a small subset of ribosomal proteins. Activation of this pathway ultimately results in eEF1A degradation, presumably clearing the GTPase center and allowing elongation to resume.

A detailed mechanistic understanding of how RNF14 and RNF25 cooperate with GCN1 to promote ubiquitination of stalled eEF1A/ribosome complexes will require biochemical reconstitution of this multilayered signaling pathway from purified components, along with structural analysis of key intermediates. Nevertheless, by integrating our data with recently published work, we can begin to build a coherent (if speculative) model (**Figure 7A**). Because a fraction of RNF25 interacts with ribosomes even in unstressed cells, we speculate that it plays an essential ’upstream’ surveillance role, perhaps by promoting RPS27A K107/K113 ubiquitination in response to pausing or stalling events. Consistent with this idea, we also detected RPS27A K107/K113 ubiquitination in untreated cells, which subsequently increased 2- fold after treatment with ternatin-4. Of note, ubiquitination of RPS27A K107/K113 was uniquely and solely dependent on RNF25 (and not RNF14). Elevated RPS27A ubiquitination requiring active translation has previously been observed in cells lacking the deubiquitinase USP16 (Montellese et al., 2020) or treated with low-dose emetine (Sinha et al., 2020), although the relevant E3 ligase was not identified in either study. We speculate that RNF25, possibly via direct ubiquitination of RPS27A K107/K113, provides one of two signaling inputs required to activate the RBR-type E3 ligase, RNF14; a potential mechanism for this requirement is described below.

GCN1 provides a second essential signaling input, most likely by recruiting and activating RNF14 near the GTPase center of elongation-stalled ribosomes. The cryo-EM structure of yeast Gcn1 bound to collided di-ribosomes suggests a potential mechanism (Pochopien et al., 2021), as the RWD binding domain of Gcn1 is positioned adjacent to the GTPase center of the leading (stalled) ribosome (**Figure 7B**). Hence, binding of this highly conserved region to the N-terminal RWD domain of RNF14 could facilitate proximity-based ubiquitination of eEF1A trapped in the GTPase center, as well as GCN1 itself (all identified GCN1 Ub sites map to the C-terminal region) and the ribosomal proteins RPL12, RPLP0, and RPLP1. Although further studies are required to define the functions of these RNF14/25- dependent ubiquitination sites, we provide evidence here for direct RNF14-mediated ubiquitination of eEF1A K385, which is essential for maximal eEF1A degradation.

The yeast Gcn1/di-ribosome cryo-EM structure also provides clues to a potential role for RNF25-dependent ubiquitination of RPS27A (aka eS31). These ubiquitination sites (K107/K113), which reside at the tip of the small ribosomal subunit ’beak’, are immediately adjacent to the RWD binding domain of Gcn1 (**Figure 7B**). This arrangement suggests a mechanism whereby RNF14 could receive two proximal signaling inputs: one provided by GCN1 binding to the RNF14 RWD domain, and a second provided by ubiquitinated K107 or K113 on RPS27A. The latter could potentially engage a conserved allosteric ubiquitin binding site in the RBR domain of RNF14. Although speculative, this model is consistent with structural and biochemical studies of other RBR-type E3 ligases (e.g., parkin and HOIP), which are autoinhibited and require allosteric activation – including by ubiquitin itself – to unveil the E2 binding and catalytic sites (Cotton and Lechtenberg, 2020). Alternatively, RNF25 could extend polyubiquitin chains from RNF14-initiated ubiquitin marks on eEF1A, analogous to the mechanism by which cullin-RING E3 ligases cooperate with the RBR E3, ARIH1 (Scott et al., 2016). We note that the RNF25 RWD domain shares the highest sequence similarity with the RWD domain of human GCN2 (EIF2AK4), whereas the RNF14 RWD domain is most similar to human IMPACT (via BLAST search), which also binds GCN1 (Pereira et al., 2005). While further studies are required to establish the mechanistic details, these observations, combined with our co-immunoprecipitation data, suggest that GCN1 may recruit RNF14, and possibly RNF25, to the GTPase center of stalled ribosomes via sequential, dynamic interactions with their RWD domains.

Historically, natural products with distinct proteotoxic mechanisms have been used to elucidate cellular stress response pathways, including the UPR (tunicamycin), mitochondrial UPR (antimycin), and ribotoxic stress response (anisomycin). A strength of our approach is the use of ternatin-4 to acutely induce ribosome stalls with trapped eEF1A, providing a robust elongation stress signal that culminates in ubiquitin-dependent eEF1A degradation. A key question for the future concerns the physiological and environmental stressors that activate the GCN1/RNF14/25 pathway in cells and organisms. One potential environmental stressor is RNA-damaging UV light, which was recently shown to induce ribosome collisions (Wu et al., 2020).

Consistent with this hypothesis, a previously published diGly proteomics dataset identified UV-induced ubiquitination sites on eEF1A, ribosomal proteins, and GCN1 (Elia et al., 2015), several of which we found to be RNF14/25 dependent and/or strongly induced by ternatin-4 treatment (**Figure S6**).

In addition to RNA damage, altered post-transcriptional RNA modifications – in particular, decreased methoxycarbonylmethylation or thiolation of the anticodon wobble uridine (U34) in certain tRNAs – could result in codon-specific stalls that locally activate GCN1/RNF14/25. Preliminary support for this idea is provided by a recent study that correlated genome-wide CRISPR knockout effects (https://depmap.org) across 485 cancer cell lines (Wainberg et al., 2021). Based on statistical analysis of gene-gene correlations (’co-essentiality’), RNF25 was placed in the same biological pathway as ELP3 and CTU2. The latter enzymes promote tRNA U34 methoxycarbonylmethylation and thiolation, respectively, and control translation elongation in developmental, homeostatic, and disease contexts (Hawer et al., 2019; Hermand, 2020). Finally, ribosome stalling at termination codons could also locally activate the GCN1/RNF14/25 pathway, leading to ubiquitination of eEF1A (bound to near-cognate aa-tRNA) or release factors (eRF1/eRF3) trapped in the GTPase center. Consistent with this scenario, overexpression of the RNF14 ortholog Itt1 promotes stop codon readthrough in yeast (Urakov et al., 2001), although the mechanism underlying this phenotype remains unclear. Analysis of expanded CRISPR knockout datasets (Depmap portal, 22Q1 release, 1070 cell lines) reveals additional strong correlations between RNF25 and the termination/splitting factors, ETF1 (aka eRF1, p = 4.5e-16) and ABCE1 (p = 1e-13), both of which are ranked among the top 100 out of 17K genes tested. RNF25 and RNF14 knockout effects are also significantly correlated across the 1070 cell lines (p = 5e-9, all p-values uncorrected), albeit to a lesser extent.

Deletion of GCN1, or even the C-terminal RWD binding region of GCN1, is lethal in mice (Yamazaki et al., 2020), whereas deletion of GCN2 causes only mild phenotypes (Zhang et al., 2002). Hence, GCN1 must have GCN2-independent functions, which have remained enigmatic despite the identification of other RWD domain-containing Gcn1 interactors, including yeast Gir2 and Yih1 (Pochopien et al., 2021; Sattlegger et al., 2004; Wout et al., 2009). We speculate that GCN1 lies at the nexus of multiple elongation stress pathways requiring RWD domain effectors – including RNF14 and RNF25 – in which pathway selection is determined locally by the status and occupancy of the GTPase center and A site of the stalled ribosome.

## Acknowledgments

Funding for this study was provided by the UCSF Invent Fund (J.T.), the National Institutes of Health (DP2 GM119139 to M.K. and P30CA082103 to the UCSF HDFCCC Laboratory for Cell Analysis Shared Resource Facility), a UCSF Genentech Fellowship (K.O.), a Department of Defense NDSEG fellowship (S.K.S.), and the Tobacco-Related Disease Research Program Postdoctoral Fellowship Awards (28FT-0014 to H.Y.W.).

## METHODS

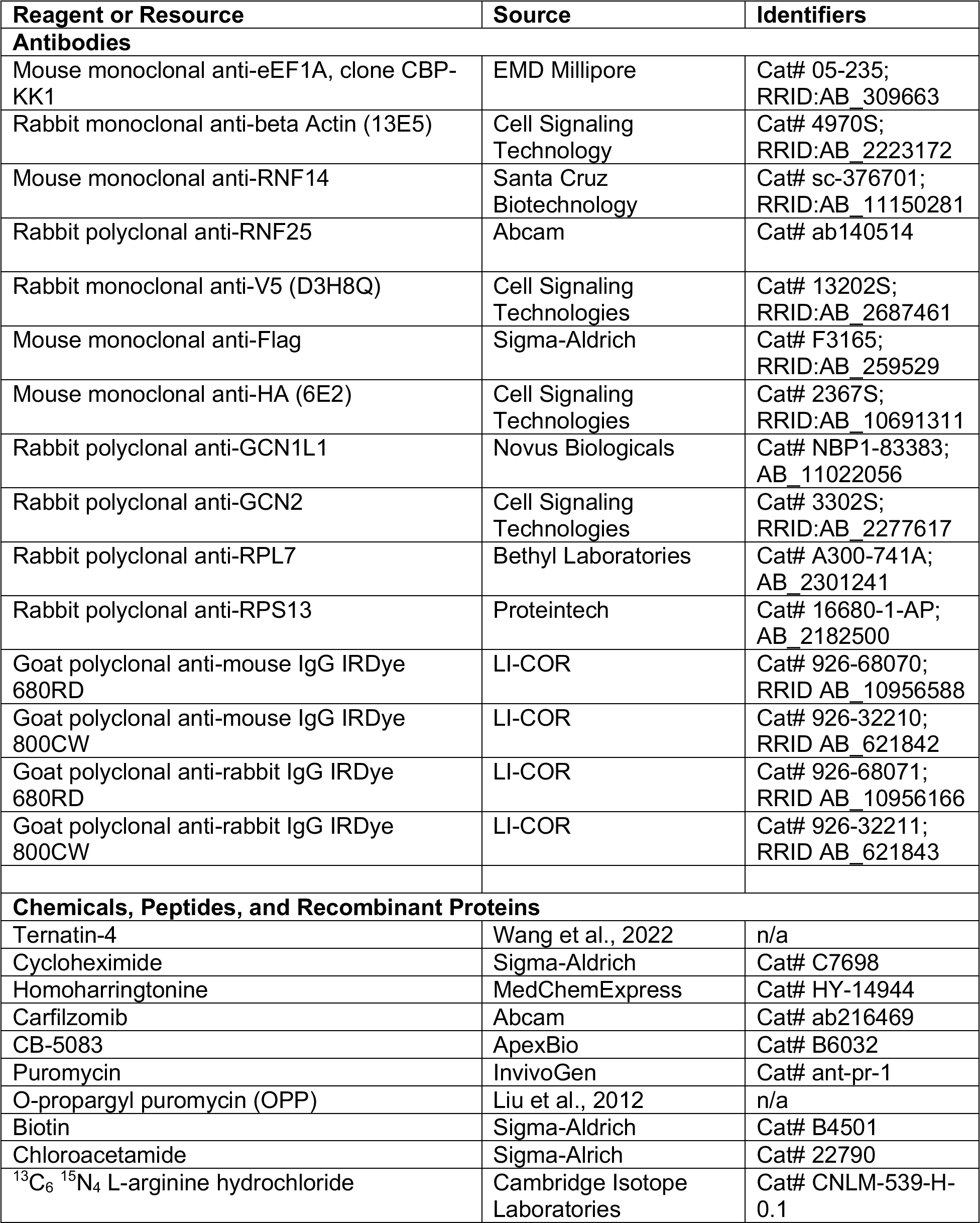
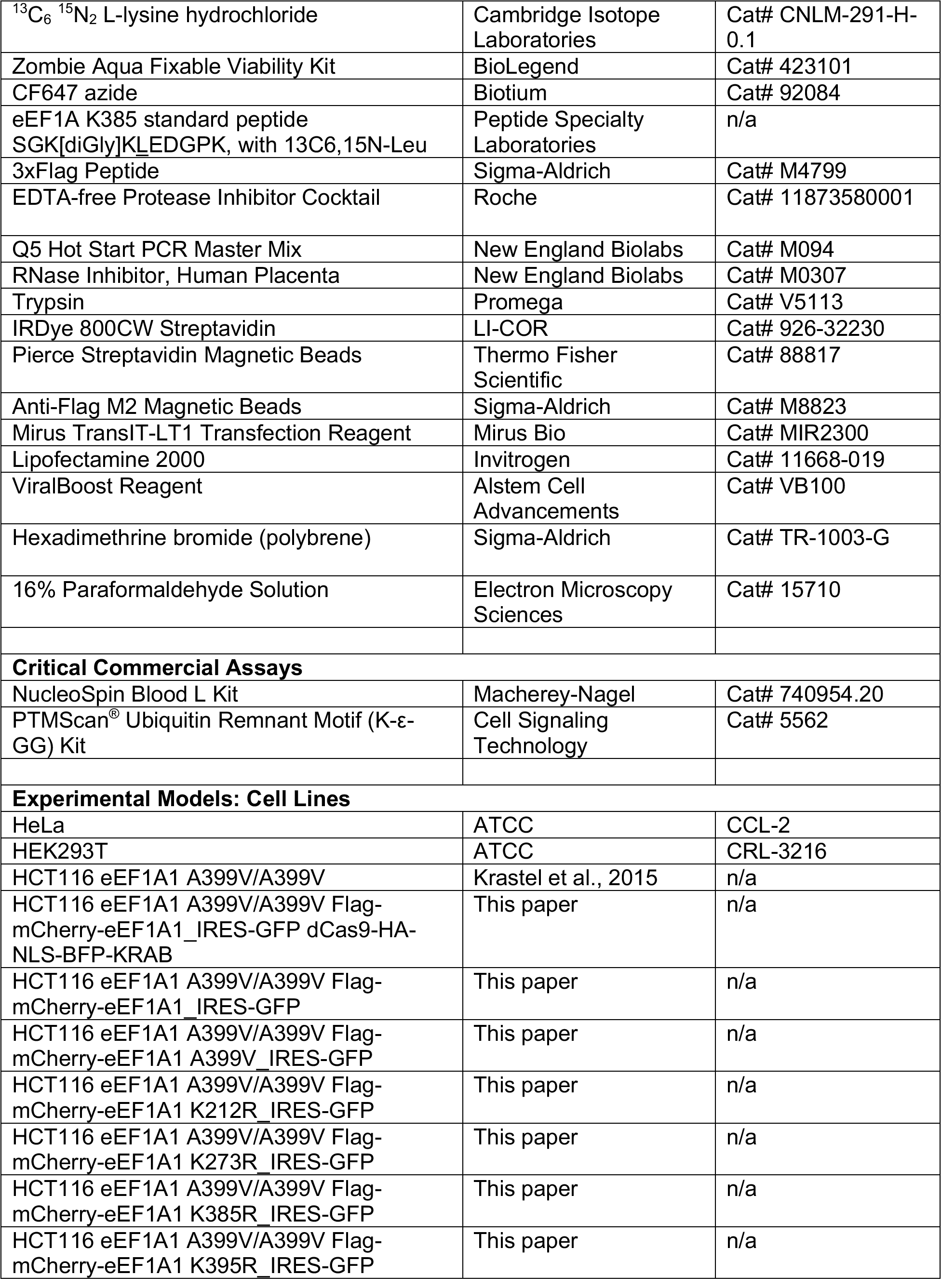
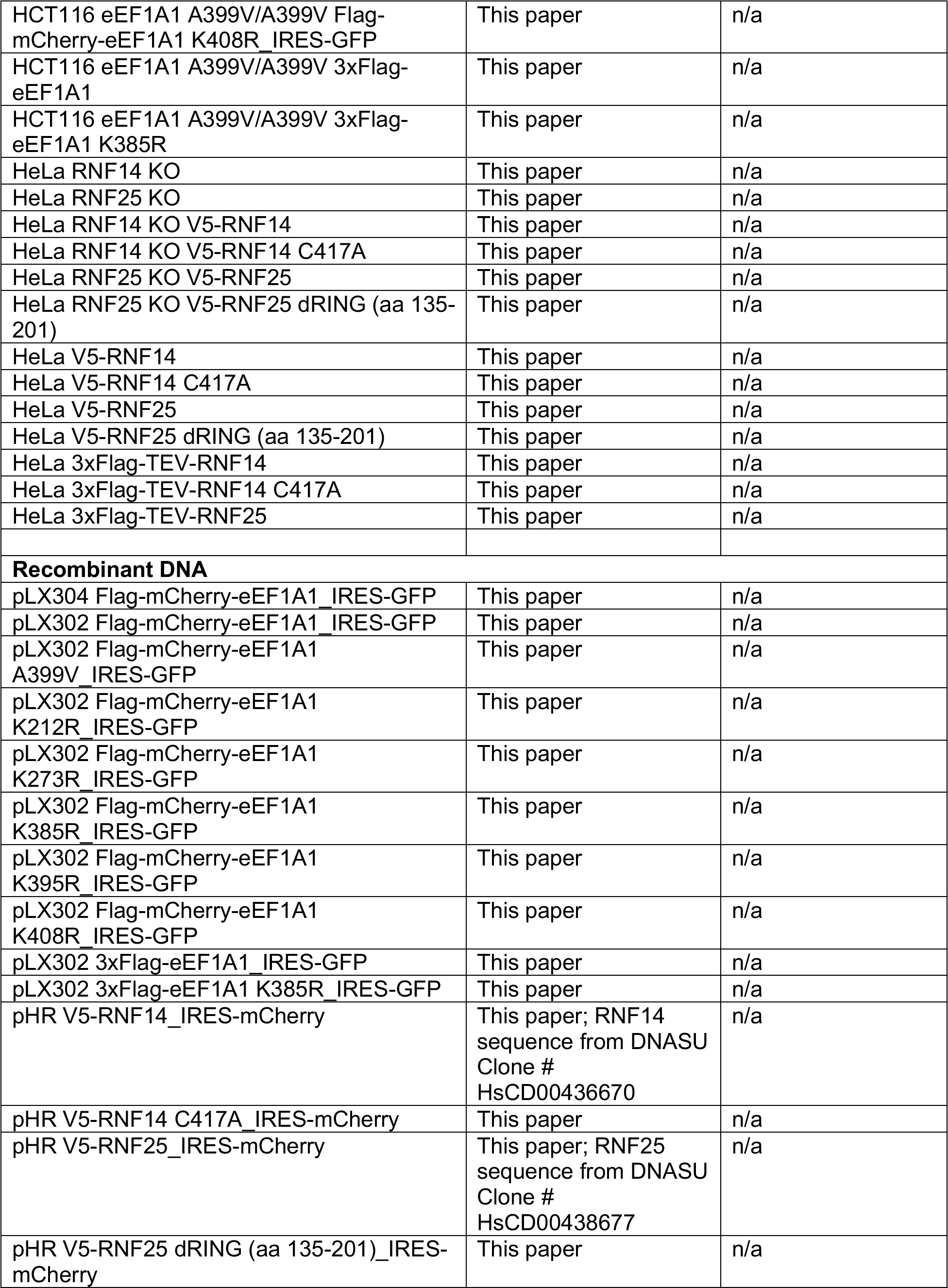
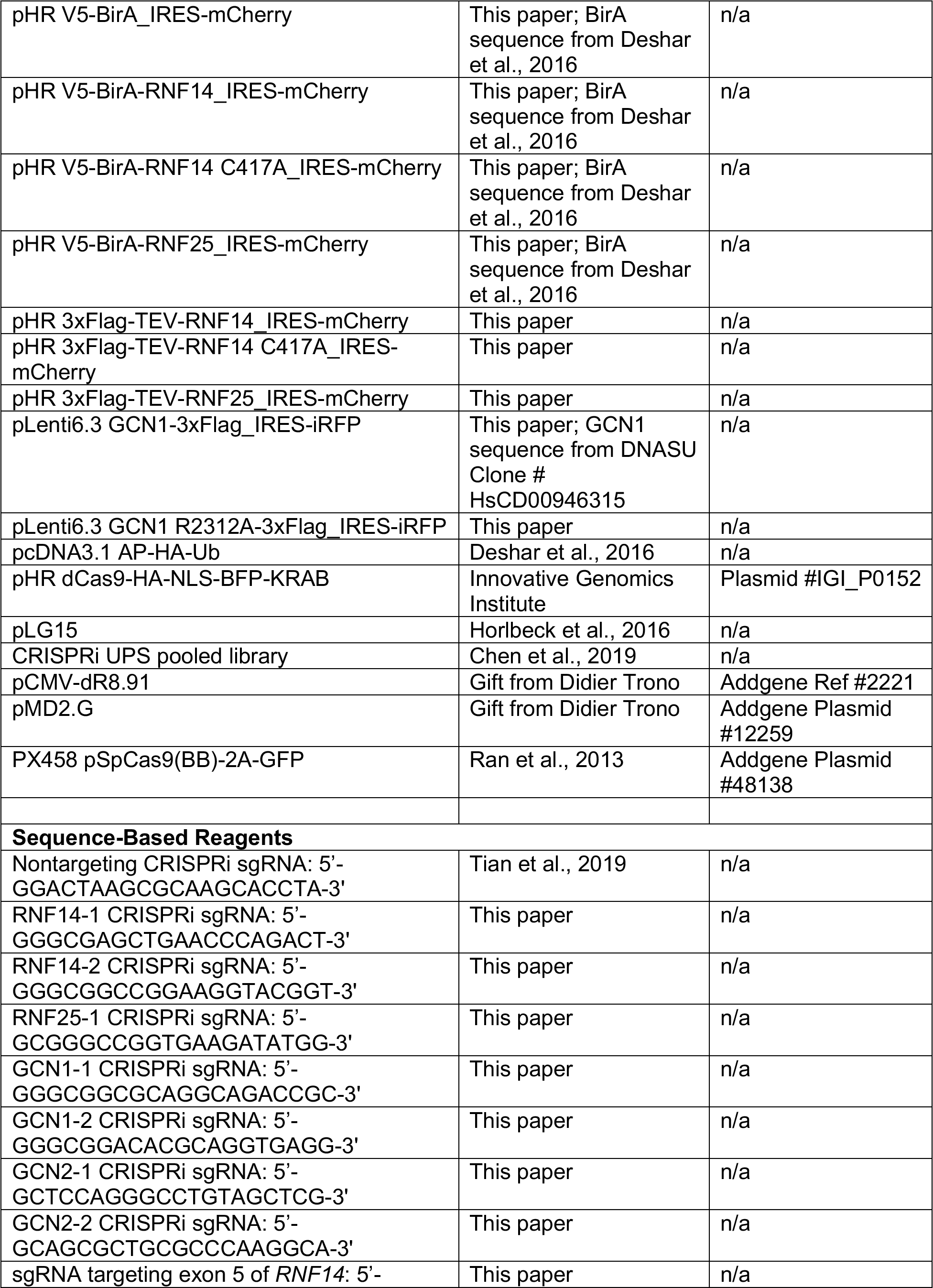
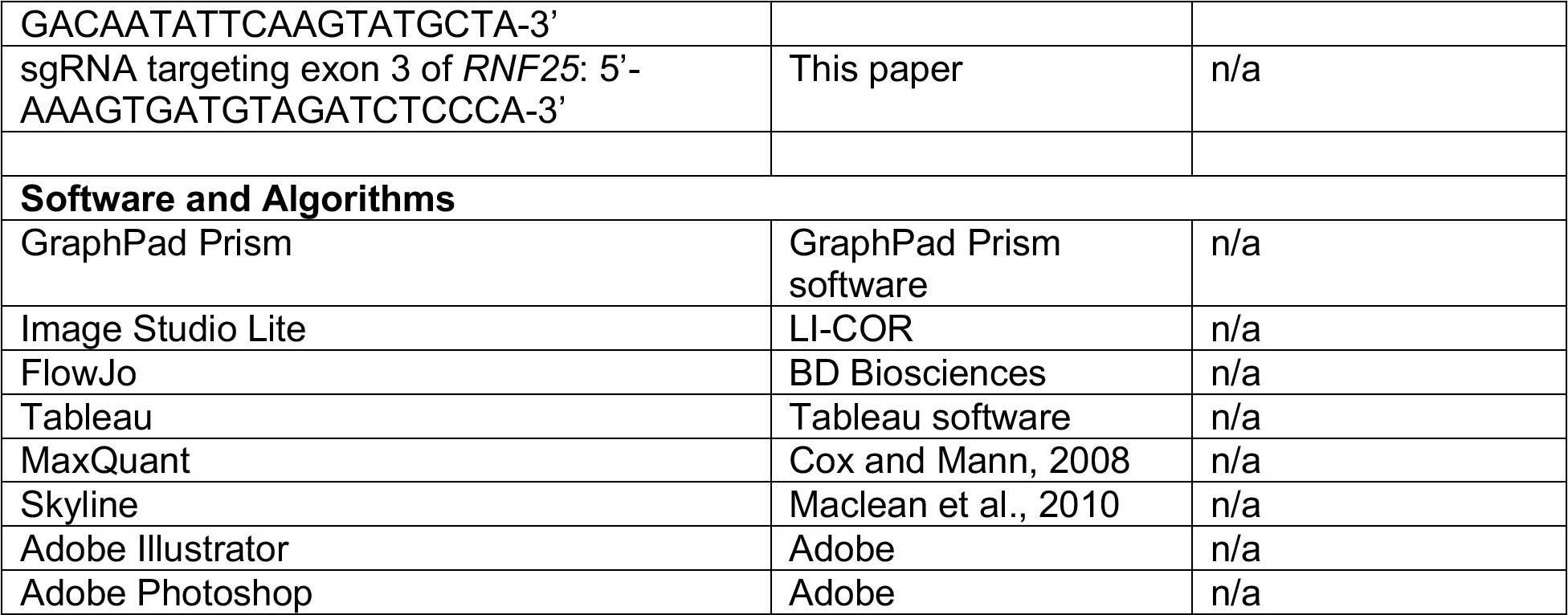
Key Resources Table

### CONTACT FOR REAGENT AND RESOURCE SHARING

Further information and requests for reagents should be directed to the lead contact, Jack Taunton.

### EXPERIMENTAL MODEL AND SUBJECT DETAILS

#### Cell lines and culture conditions

HEK293T and HeLa cells were maintained in Dulbecco’s Modified Eagle’s Medium (DMEM) supplemented with 10% fetal bovine serum (FBS), 100 U/mL penicillin, and 100 µg/mL streptomycin. HCT116 eEF1A1 A399V/A399V cells were originally described in Krastel et al., (2015), and were maintained in McCoy’s media supplemented with 10% FBS, 100 U/mL penicillin, and 100 µg/mL streptomycin. Antibiotics were omitted for all transfection-based experiments. Cells were grown in a humidified incubator at 37 °C with a 5% CO2 atmosphere. Unless indicated otherwise, all treatment conditions utilized a vehicle control, 50 nM ternatin-4, 50 µg/mL cycloheximide, 2 µg/mL homoharringtonine, 500 nM carfilzomib or 2.5 µM CB-5083 as described in figure legends.

### METHOD DETAILS

#### Plasmid Generation

Plasmids encoding eEF1A1, RNF14, RNF25, or GCN1, and the indicated epitope tags and IRES sequences, were constructed by Gibson cloning. Point mutations and the RNF25 RING deletion (residues 135-201) were created by site-directed mutagenesis. All eEF1A1-encoding constructs were expressed from a pLX302 vector, except the mCherry-eEF1A1 reporter used for the CRISPRi screen, which was expressed from a pLX304 vector. RNF14 and RNF25- encoding constructs were expressed from a pHR vector. GCN1 was expressed in a modified pLenti6.3 vector with the SV40 promoter and blasticidin resistance marker removed. Plasmids encoding individual sgRNAs were constructed by ligating complementary oligonucleotides (sequences indicated above in **Sequence-Based Reagents**) into the BlpI and BstXI restriction enzyme sites of pLG15 (CRISPRi) or the BbsI restriction enzyme sites of PX458 (CRISPR KO). All plasmids were verified by Sanger sequencing.

#### Lentivirus and stable cell line generation

Lentivirus was generated by transfecting HEK293T cells growing at approximately 60% confluency in 6-well dishes with lipid complexes containing 1.5 µg of a construct of interest, 1.35 µg pCMV-dR8.91, 165 ng pMD2-G, and 7.5 µL Mirus TransIT-LT1 diluted in OPTI-Mem. Alstem ViralBoost was added to cells after addition of the transfection mix. Two days post-transfection, an additional 1.5 mL complete DMEM was added to cells. The viral supernatant was collected on the third day and filtered through 0.45 µm sterile SFCA syringe filters (Thermo Scientific).

Supernatant was used immediately or stored at -20 °C.

For stable cell line generation, media containing 8 µg/mL polybrene was added to cells, lentivirus was added, and cells were incubated overnight. Lentivirus was removed from cells, and cells were expanded until cell sorting using a BD FACS Aria II. Transduced cells were typically sorted to at least 95% purity.

#### CRISPR knockout cell lines

CRISPR knockout cells were generated as previously described (Ran et al., 2013). sgRNA sequences targeting exon 5 of RNF14 and exon 3 of RNF25 were identified using the Broad CRISPR design tool (crispr.mit.edu/v1) and ligated into the PX458 (pSpCas9(BB)-2A-GFP) plasmid. HeLa cells were transfected using Lipofectamine 2000 and sorted for GFP positivity after 3 days. Eight days post-transfection, cells were plated into a 96-well plate at 0.5 cells per well. After appearance of visible colonies (approximately 2 weeks), wells containing individual colonies were collected by trypsinization and expanded to 6-well dishes. Clones were screened for RNF14/RNF25 expression by Western blotting and Sanger sequencing prior to selection for experiments.

#### Immunoblot analysis

Cells were washed once with ice-cold PBS and stored at -80 °C or immediately lysed using lysis buffer (50 mM HEPES pH 7.4, 150 mM NaCl, 1% NP-40, 2x Roche EDTA-free protease inhibitors). Lysates were collected by scraping and were cleared by centrifugation at 16,100 *g* for 10 minutes at 4 °C. Total protein was quantified using the Bradford method and normalized prior to electrophoresis using hand-cast 7.5% polyacrylamide gels. Proteins were transferred to 0.45 µm nitrocellulose membranes (Bio-Rad) using a Bio-Rad Criterion transfer system. Membranes were blocked for 1 hour at room temperature using blocking buffer (5% BSA, 0.1% sodium azide in TBS-T). Membranes were incubated with primary antibodies diluted in blocking buffer for 1 hour at room temperature or overnight at 4 °C. Membranes were rinsed with TBS-T (3 x 5 minutes at room temperature) and incubated with secondary antibodies diluted in blocking buffer for 1 hour at room temperature. Membranes were rinsed with TBS-T (3 x 5 minutes at room temperature) and were imaged using a Li-Cor Odyssey system.

#### Protein synthesis measurements

Cells were grown to 70% confluency in 12-well plates and treated as indicated (ternatin-4 or vehicle; 20 h). Following treatment, O-propargyl puromycin (OPP) was added to all wells at a final concentration of 30 µM, and cells were incubated for an additional hour. Cells were collected by trypsinization and transferred to a 96-well V-bottom plate for the remainder of the experiment. All centrifugation steps were performed at 2,100 *g* for 3 minutes at 4°C, and all washes and incubations were done with 200 µL buffer unless indicated otherwise. Cells were washed once with cold PBS and then stained with 100 µL Zombie Aqua (BioLegend; 1:1000 in PBS) for 30 min at RT. Incubations from this step onwards were performed in the dark. Cells were washed once (2% FBS in PBS) and fixed with 4% paraformaldehyde (PFA) in PBS for 15 min at 4 °C. Cells were washed once (2% FBS in PBS) and incubated in permeabilization buffer (3% FBS, 0.1% saponin in PBS) for 5 min at RT. Click chemistry mix was prepared by adding the following components to final concentrations in the order listed: 50 mM HEPES pH 7.5, 150 mM NaCl, 400 µM TCEP, 250 µM TBTA, 5 µM CF647 azide (Biotium), 200 µM CuSO4. Cells were resuspended in 25 µL permeabilization buffer, 100 µL click chemistry mix was added, and cells were incubated overnight at RT. Cells were washed twice with permeabilization buffer and twice with FACS buffer (2% FBS, 100 U/mL penicillin, 100 µg/mL streptomycin, and 2 mM EDTA in PBS lacking Ca^+2^/Mg^+2^). Samples were analyzed on a Thermo Attune NxT (see below). Data was processed in FlowJo (BD) with a gating hierarchy as follows: debris were excluded (FSC-H vs. SSC-A), doublets were excluded (FSC-H vs. FSC-W), and dead cells were excluded on the basis of Zombie Aqua positivity. Mean fluorescent intensity (MFI) was calculated for each sample and normalized to vehicle control samples.

#### Flow cytometry

Cells were harvested by trypsinization, transferred to 96-well V-bottom plates, and centrifuged at 2,100 *g* for 3 minutes at 4 °C. Cells were resuspended in FACS buffer (2% FBS, 100 U/mL penicillin, 100 µg/mL streptomycin, and 2 mM EDTA in PBS lacking Ca^+2^/Mg^+2^) and analyzed using a Thermo Attune NxT equipped with 405 nm, 488 nm, 561 nm, and 637 nm lasers. Single cells were analyzed using FlowJo (BD) software.

#### CRISPRi screen

CRISPRi screening cells were generated by transducing ternatin-resistant HCT116 eEF1A1 A399V/A399V cells with lentivirus encoding dCas9-BFP-KRAB. Cells were sorted twice for BFP positivity. The resulting dCas9-containing cells were transduced with the eEF1A FACS reporter (pLX304 Flag-mCherry-eEF1A1_IRES-AcGFP) and were sorted twice for GFP positivity.

Throughout the screen, a minimum of 1000x representation was maintained for sgRNA elements. A sgRNA library targeting elements of the ubiquitin proteasome system (Chen et al, 2019; 9,564 sgRNAs total) was utilized. Several sgRNAs targeting additional eEF1A or RQC-related genes (*EF1A1*, *EF1A2*, *HBS1L*, *PELO*, *NEMF*, *TCF25*, *PUM2*, *PCBP1*) were individually cloned (5 sgRNAs/gene) and added to the library. Lentivirus was generated by transfecting two 15 cm dishes of HEK293T cells each with 9 µg of library, 8 µg of pCMV-dR8.91, and 1 µg of pMD2-G per dish using Mirus Trans-IT LT1 and Alstem ViralBoost. Viral supernatant was collected after two days and filtered. Six 15 cm dishes of CRISPRi screening cells were transduced with all freshly harvested virus, yielding a transduction efficiency of approximately 55% (measured by FACS two days post-transduction). Puromycin selection for sgRNA-containing cells began three days post-transduction and was conducted for 48 hours, with daily replenishment of media containing 2 µg/mL puromycin. Six days post-transduction, cells were plated into two sets of 8 15 cm dishes (96 million cells per set) for treatment and cell sorting the following day.

Cells at approximately 70% confluency were left untreated or treated for 8 hours with 50 nM ternatin-4. Cells were harvested by trypsinization, resuspended in FACS buffer (2% FBS, 100 U/mL penicillin, 100 µg/mL streptomycin, and 2 mM EDTA in PBS lacking Ca^+2^/Mg^+2^), and sorted on a BD FACS Aria II equipped with BD FACSDiva software. Single, BFP positive cells were separated into high, middle, and low thirds based on the calculated ratio of mCherry to GFP for each cell, with approximately 9 million cells collected for each population. Sorted cell populations were washed once with PBS and stored at -80°C until genomic DNA extraction.

Sorted samples were prepared for next-generation sequencing as previously described (Tian et al., 2019). Briefly, genomic DNA was isolated from samples using the Machery-Nagel Blood Midi kit (catalog number 740954.20) according to the manufacturer’s instructions. TSS (transcription start site)-specific sgRNA sequences were PCR amplified using all genomic DNA, indexed PCR primers, and 2x Q5 Hot Start PCR Master Mix (NEB M094). In total, approximately 40-55 60 µL PCR reactions were prepared for each sample. PCR products were size-selected using SPRI-select beads to remove PCR primers and genomic DNA, and samples were sequenced using an Illumina HiSeq 4000.

#### Individual sgRNA knockdown and rescue

Lentivirus was generated for plasmids encoding individual sgRNAs. CRISPRi reporter cells (ternatin-resistant HCT116 eEF1A1 A399V/A399V expressing Flag-mCherry-eEF1A1_IRES- AcGFP and dCas9-BFP-KRAB) were transduced, and were selected three days post-transduction with 2 µg/mL puromycin for 48 hours.

Typically, sgRNA knockdown cells were plated into 12-well dishes for experiments 6 days post-transduction. The next day, cells were treated and analyzed as indicated in figure legends. For GCN1 rescue experiments, puromycin selected, sgRNA-containing cells were transfected in 6- well dishes with 1 µg of the indicated GCN1 constructs (1 well/construct) using the manufacturer’s protocol for Lipofectamine 2000 (Thermo Fisher). The next day, cells were distributed into 12-well dishes for flow cytometry analysis or 6-well dishes for immunoblotting analysis, and incubated overnight. Cells were treated and analyzed as indicated in the figure legends.

#### DiGly peptide enrichment

Cells were grown in light (Arg0/Lys0) or heavy (Arg10/Lys8) SILAC DMEM (Thermo Scientific) supplemented with 10% dialyzed FBS, 100 U/mL penicillin,100 µg/mL streptomycin, 80 µM lysine0 or lysine8, and 40 µM arginine0 or arginine10 for 12 days prior to indicated treatments. Typically, 5-6 15 cm dishes were used per label and treatment condition, yielding approximately 10-15 mg total protein. A label swap was performed as a biological replicate for all experimental conditions. Cells were harvested at room temperature by quickly aspirating media from dishes, rinsing once with ice-cold PBS, and scraping cells into cell lysis buffer (20 mM HEPES pH 8.0, 9 M urea, 1 mM sodium orthovanadate, 2.5 mM sodium pyrophosphate, 1 mM beta-glycerophosphate). Lysates were sonicated using three 15-second bursts from a microtip sonicator, and lysates were clarified by centrifugation at 19,000 *g* for 15 minutes at RT. Cleared lysates were quantified using the BCA method, and equal amounts of heavy and light samples were pooled (20-30 mg total). Samples were reduced with 4.5 mM DTT for 30 minutes at 55 °C. Samples were allowed to cool to room temperature and alkylated with 10.2 mM chloroacetamide for 30 minutes at RT. Lysates were diluted 4-fold with 20 mM HEPES pH 8.0 to a final concentration of approximately 2 M urea, sequencing-grade trypsin (Promega V5113) was added at a 1:300 w/w ratio, and samples were digested by rotating overnight at RT. The next day (approximately 18 hours later), trypsin digestion was stopped by addition of TFA to 1% final concentration. Samples were incubated on ice for 15 minutes and centrifuged at 1,780 *g* for 15 minutes at RT. Peptides were desalted using Waters 500 mg C18 SepPaks (WAT036945).

SepPak cartridges were conditioned with 7 mL acetonitrile and equilibrated with 3 sequential washes of 1.4 mL, 4.2 mL, and 8.4 mL Solvent A (0.1% TFA in water) prior to loading of the clarified peptide solution. Peptides were loaded onto columns by gravity flow, and columns were washed 3 times using 1.4 mL, 7 mL, and 8.4 mL Solvent A. Peptides were eluted by 3 sequential additions of 2.8 mL solvent B (0.1% TFA, 40% acetonitrile in water). Samples were frozen and lyophilized for a minimum of 48 hours prior to immunoprecipitation of diGlycine- containing peptides.

Ubiquitin remnant-containing peptides were isolated using the Cell Signaling Technologies PTMScan Ubiquitin Remnant Motif Kit (CST #5562). Lyophilized peptides were collected by centrifugation at 2,000 *g* for 5 minutes at RT and dissolved in 1.4 mL ice-cold Immunoaffinity Purification (IAP) buffer (50 mM MOPS pH 7.2, 10 mM disodium phosphate, 50 mM NaCl). The peptide solution was cleared by centrifugation at 10,000 *g* for 5 minutes at 4 °C. The peptide supernatant was added to a tube containing immunoaffinity beads pre-washed 4x with 1 mL of cold PBS. All centrifugation steps with antibody beads were done at 2,000 *g* for 30 seconds at 4 °C. Immunoprecipitations were performed using a predetermined ratio of 10 mg input protein per 10 µL of antibody bead slurry (Udeshi et al., 2013). Samples were rotated for 2 hours at 4 °C, and subsequently, peptide supernatant was removed from beads. Beads were washed twice with 1 mL ice-cold IAP buffer and 3 times with 1 mL ice-cold HPLC water. Peptides were eluted from beads with two consecutive incubations of 55 µL and 50 µL 0.15% TFA for 10 minutes at RT with gentle agitation every 2-3 minutes. Eluants were desalted using Agilent OMIX C18 tips (A57003100), dried by vacuum concentration, and stored at -80 °C until analysis.

#### LC-MS/MS data acquisition (DDA)

One half of each sample was resuspended in 10 µL 0.1% FA or 10 µL 0.1% FA containing 2.5 nM eEF1A K385 standard peptide (SGK[diGly]K<u>L</u>EDGPK, with ^13^C6,^15^N-Leu). One quarter of each reconstituted sample (2.5 µL; 12.5% of total sample) was analyzed on an Orbitrap Fusion Lumos (Thermo Scientific) equipped with an ACQUITY M-Class UPLC (Waters) and an EASY-Spray C18 column (Thermo Scientific #ES800; 75 μm x 15 cm, 3 μm particle size, 100 Å pore size). Solvent A was 0.1% FA in water, and Solvent B was 0.1% FA in acetonitrile. Peptides were loaded onto the warmed column (45 °C), and equilibrated with 0% B for 13 min using a flow rate of 600 nL/min, followed by 0-30% B over 120 min at 300 nL/min, 30-80% B over 20 min, 80% B for 5 min, 80%-0% B over 2 min, and 0% B for 10 min with the flow rate changed back to 600 nL/min. The entire method ran for a total of 170 min. Mass spectra were acquired in data dependent mode. MS1 scans were collected in the orbitrap at a resolution of 120,000, a scan range of 375-1500 m/z, an automatic gain control (AGC) target of 4E5, and a maximum injection time of 50 ms. Ions with a peptide-like isotopic distribution (MIPS set to “peptide”) that exceeded an intensity threshold of 2E4 and contained a charge between 2 and 7 were selected for HCD fragmentation. MS2 spectra of HCD-fragmented peptides were collected using a 1.6 m/z isolation window and an HCD collision energy of 30%. Fragment ions were measured in the orbitrap at a resolution of 30,000 and were collected with an AGC target of 5E4 and a maximum injection time of 100 ms. Peptides selected for fragmentation were dynamically excluded for the following 30 seconds using a 10-ppm window. The maximum duty cycle was set to 3 s.

#### PRM analysis of eEF1A K385 ubiquitination

Sample reconstitution and liquid chromatography proceeded identically to DDA preparations (see above). For each cycle of the PRM method, an MS1 scan was first collected with a scan range of 360-1300 m/z, a resolution of 60,000, an automatic gain control (AGC) target of 4E5, and a maximum injection time of 50 ms. Subsequently, targeted MS2 scans were collected for heavy, light, and standard (medium) eEF1A K385 peptides (z = 3; m/z = 399.5614, 391.5472, and 393.8862, respectively). MS2 precursors were fragmented by HCD using a collision energy of 30%. MS2 scans utilized an isolation window of 0.7 m/z, a resolution of 15,000, an AGC target of 1E5, and a maximum injection time of 150 ms. The entire scan cycle was repeated over the entire gradient. Data was analyzed with Skyline using a reference library constructed from previous DDA runs with the same samples. The six most highly ranked (most intense) b or y ions were chosen for generation of extracted ion chromatograms.

#### Proximity-based ubiquitination assay

Ternatin-resistant HCT116 cells (eEF1A1 A399V/A399V) were cultured in DMEM, which does not contain biotin, supplemented with 10% fetal bovine serum. The assay was based on Deshar et al., (2016), with some modifications. Cells were grown to 70% confluency in 10 cm dishes and were co-transfected with 1.8 µg AP-HA-Ub (pcDNA3.1) and 8.8 µg V5-BirA or V5-BirA-RNF (pHR) constructs using the manufacturer’s protocol for Lipofectamine 2000. The next day, cells were treated with 50 µM biotin and with DMSO or 50 nM ternatin-4 for 4 h. Following treatment, cells were washed once with ice-cold PBS and collected by scraping into PBS. Cells were centrifuged at 1000 *g* for 3 minutes at 4 °C, and cell pellets were frozen in liquid nitrogen and stored at -80 °C or immediately lysed. Cells were lysed in lysis buffer (50 mM HEPES pH 7.4, 150 mM NaCl, 1% NP-40, 2x Roche EDTA-free protease inhibitors, and 50 mM chloroacetamide) and lysate was cleared by centrifugation at 16,100 *g* for 10 minutes at 4 °C. SDS was added to supernatants to 1% final concentration, and samples were boiled for 10 minutes. Denatured lysates were quantified by the Bradford method, normalized, and diluted 10- fold in lysis buffer. Diluted lysates were filtered through 0.45 µm SFCA syringe filters (Thermo Scientific) and were added to tubes containing streptavidin magnetic beads (Pierce; 40 µL beads per 1 mg protein). Samples were rotated for 2 hours at 4 °C. Supernatants were removed from beads, and beads were washed twice with wash buffer 1 (50 mM HEPES pH 7.4, 150 mM NaCl, 1% NP-40, 0.1% SDS), and twice with wash buffer 2 (50 mM HEPES pH 7.4, 500 mM NaCl, 1% NP-40). Bound proteins were eluted by boiling beads for 10 minutes in elution buffer (50 mM HEPES pH 7.4, 150 mM NaCl, 1% NP-40, 5 mM biotin, and 1x SDS loading buffer). 10 µg of input protein (typically < 3% total input protein) and 40% of eluants were loaded on 7.5% gels and analyzed by immunoblotting.

#### Co-immunoprecipitations

For GCN1-3xFlag immunoprecipitations, HEK293T cells growing in a 15 cm dish at approximately 70% confluency were transfected with 8 µg plasmid (pLenti6.3) using Lipofectamine 2000. The next day, cells were split between two 15 cm dishes. Cells were treated approximately 48 hours post-transfection. For 3xFlag-RNF14 or RNF25 immunoprecipitations, HeLa cells stably expressing the indicated construct were were grown to approximately 70% confluency and treated in 10 cm dishes.

Cells were harvested by scraping into ice-cold PBS and centrifuged at 1000 *g* for 3 minutes at 4 °C. Unless otherwise indicated, cells were crosslinked as described by Shi et al., (2017), as follows: cells were resuspended in 1 mL PBS containing 0.1% PFA (Electron Microscopy Sciences) and were rotated for 10 minutes at room temperature; crosslinking was stopped by addition of 100 µL quench solution (2.5 M glycine, 25 mM Tris pH 7.4), and cells were centrifuged at 1000 *g* for 3 minutes at 4 °C. Cells were lysed in lysis buffer (50 mM HEPES pH 7.4, 100 mM KOAc, 5 mM Mg(OAc)2, 0.1% NP-40, 40 U/mL RNAse inhibitor (NEB), 2x Roche EDTA-free protease inhibitors, and 50 mM chloroacetamide) for 5 minutes on ice, and lysates were cleared by centrifugation at 15,000 *g* for 10 minutes at 4 °C. Lysates were quantified by the Bradford method and normalized. Samples were added to tubes containing Flag M2 magnetic beads (Sigma) and rotated for 1 hour at 4 °C. For GCN1-3xFlag IPs, 40 µL bead slurry was used per 1 mg protein, and for 3xFlag-RNF14 and RNF25 IPs, 20 µL bead slurry was used per 1 mg protein. After binding, supernatants were removed and beads were washed 3 times using wash buffer (50 mM HEPES pH 7.4, 100 mM KOAc, 5 mM Mg(OAc)2, 0.1% NP-40). Two sequential elutions were performed by incubating 200 µg/mL 3xFlag peptide in wash buffer with rotation for 30 minutes at 4°C. 10 µg input protein (typically < 2% total input protein) and 40% of IP eluates were loaded onto 4-12% Bis-Tris gradient gels (Invitrogen) and analyzed by immunoblotting.

### QUANTIFICATION AND STATISTICAL ANALYSIS

#### Quantification and normalization

Immunoblots were visualized using a Li-Cor Odyssey near-IR system. The resulting images were quantified using Image Studio Lite (Li-Cor). Where indicated, intensities were normalized to vehicle controls. Flow cytometry data was quantified using the mean fluorescent intensity (MFI; protein synthesis measurements), or mean or median mCherry/GFP ratio (eEF1A degradation reporter), across a single cell population. MFI values were normalized to the vehicle control for an indicated cell line. To ensure reproducibility, three independent replicates were conducted with similar results for all immunoblotting and flow cytometry experiments.

#### CRISPRi screen analysis

Sequencing data was analyzed as previously described (Kampmann et al., 2013; Tian et al., 2019). Briefly, raw sequencing data was aligned to the reference library sequences using Bowtie, and the number of reads for each sgRNA were counted. Sequencing counts were normalized to the total number of counts within each sequenced sample. Phenotype scores for each sgRNA were calculated as the log2 ratio of high mCherry/GFP sequencing counts divided by low mCherry/GFP counts. An epsilon value was calculated by averaging the three most extreme phenotype scores for each transcription start site (TSS). A p value for each TSS was calculated using the Mann-Whitney U test against nontargeting controls. A gene score was calculated for each TSS as the product of the epsilon score and the -log10(p value). Epsilon values, p values, and products for all genes targeted in the library are included in **Supplementary Table 1**.

#### Mass spectrometric site identification and quantification

The raw data were searched against the human Uniprot database (73,651 entries) using MaxQuant (version 1.6.7.0) with a list of common laboratory contaminants added. ‘Arg10’ and ‘Lys8’ were selected as heavy labels. The digestion enzyme was set to trypsin, a maximum of three missed cleavages were allowed, and the minimum peptide length was set to 7. Cysteine carbamidomethylation was specified as a fixed modification, and N-terminal protein acetylation and methionine oxidation were set as variable modifications. Internal lysine diGlycine modifications, and lysine diGlycine modifications at the protein C-terminus were additionally set as variable modifications. The remainder of the search parameters were left at default settings. Peptide requantification was enabled for calculation of H/L SILAC ratios. DiGly site identifications and normalized SILAC ratios were obtained from the MaxQuant sites file. SILAC ratios were inverted for label swap experiments. Site values were averaged between biological replicates, log2-transformed, and reported in **Supplementary Table 2**.

## Author Contributions

K.O. designed and performed experiments, and analyzed and interpreted data. J.D.C. designed experiments and developed the fluorescent reporter assay. S.K.S. assisted with planning and performing the CRISRPi screen and analyzed screen data. T.Y. collected mass spectrometry data and assisted with mass spectrometry data analysis. H.Y.W. synthesized key reagents.

M.K. assisted with planning the CRISPRi screen and interpreted data. J.T. supervised the study. J.D.C., K.O., and J.T. conceived the study. J.T. and K.O. wrote the manuscript with input from all authors.

---

**Figure S1.**
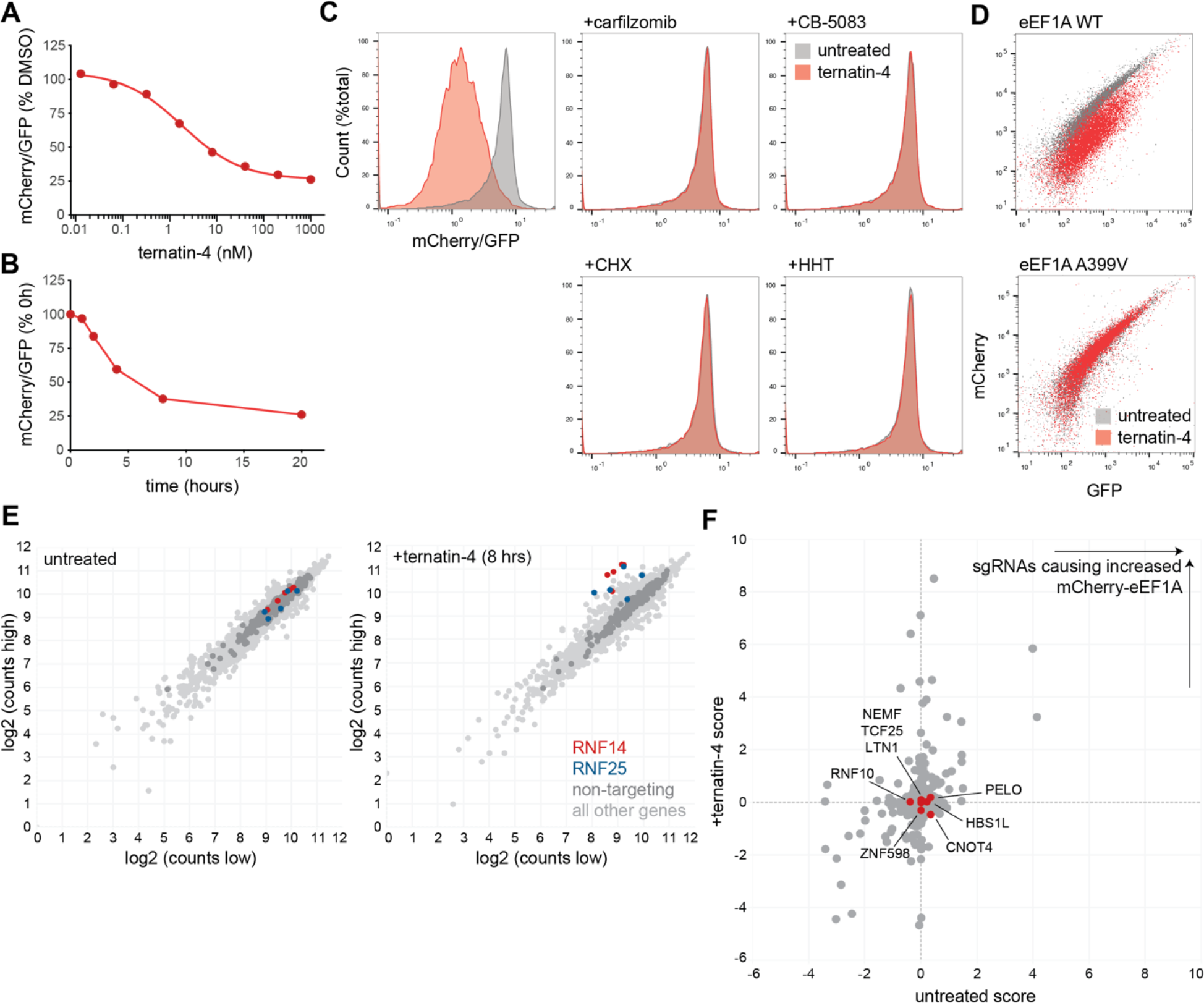
A fluorescent reporter enables CRISPRi screening for ternatin-induced eEF1A degradation. Related to Figure 2. (**A**) Dose-dependent mCherry-eEF1A reporter degradation. Ternatin-resistant HCT116 cells stably expressing the Flag-mCherry-eEF1A1_IRES-GFP reporter and dCas9-BFP-KRAB were treated with 50 nM ternatin-4 for 8 h and analyzed by flow cytometry. (**B**) Time dependence of mCherry-eEF1A reporter degradation. Cells as in (**A**) were treated with 50 nM ternatin-4 for the indicated times and analyzed by flow cytometry. (**C**) Cells in each panel were treated for 8 h ± ternatin-4 (50 nM), along with carfilzomib (500 nM), CB-5083 (2.5 μM), CHX (50 ug/mL), or HHT (2 μg/mL, 20 min pretreatment), as indicated. (**D**) Cells expressing WT or A399V mCherry-eEF1A were treated for 8 h ± ternatin-4 (50 nM) and analyzed by flow cytometry. (**E**) sgRNA sequencing counts for high and low fluorescence cell populations (± ternatin-4), highlighting RNF14 and RNF25 sgRNAs. (**F**) CRISPRi scores from Figure 2C are shown in red for HBS1L, PELO, LTN1, ZNF598, NEMF, RNF10, and CNOT4.

**Figure S2.**
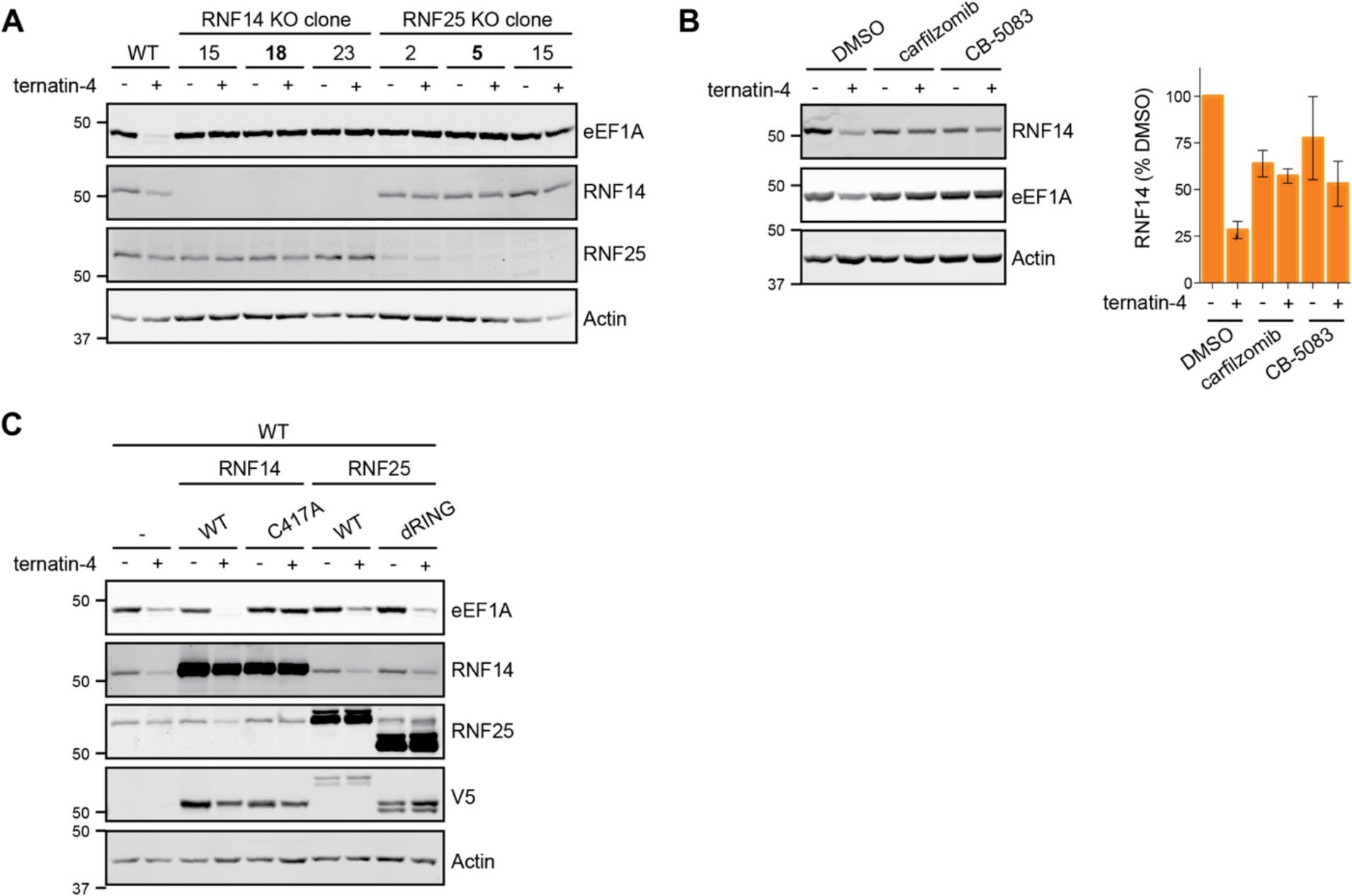
RNF14 and RNF25 are required for ternatin-induced EF1A degradation. Related to. Figure 3. (**A**) Knockout of RNF14 or RNF25 prevents ternatin-induced eEF1A degradation across multiple HeLa cell clones. CRISPR KO cell lines were constructed targeting RNF14 exon 5 and RNF25 exon 3. Cells were treated with ternatin-4 (50 nM) for 20 h and analyzed by immunoblotting. Clones 18 (RNF14) and 5 (RNF25) were utilized for further experiments. (**B**) Ternatin-induced RNF14 degradation is prevented by proteasome and p97 inhibitors. Cells were treated for 20 h as indicated. Percent RNF14 remaining in each condition was normalized to the DMSO-treated cells, and quantification is shown for n=3 experiments (mean ± sd). (**C**) Overexpression of V5-tagged RNF14, but not RNF25, increases eEF1A degradation in wild type cells. Stable cell lines were generated as in Figure 3C and treated for 20 h with ternatin-4 (50 nM).

**Figure S3.**
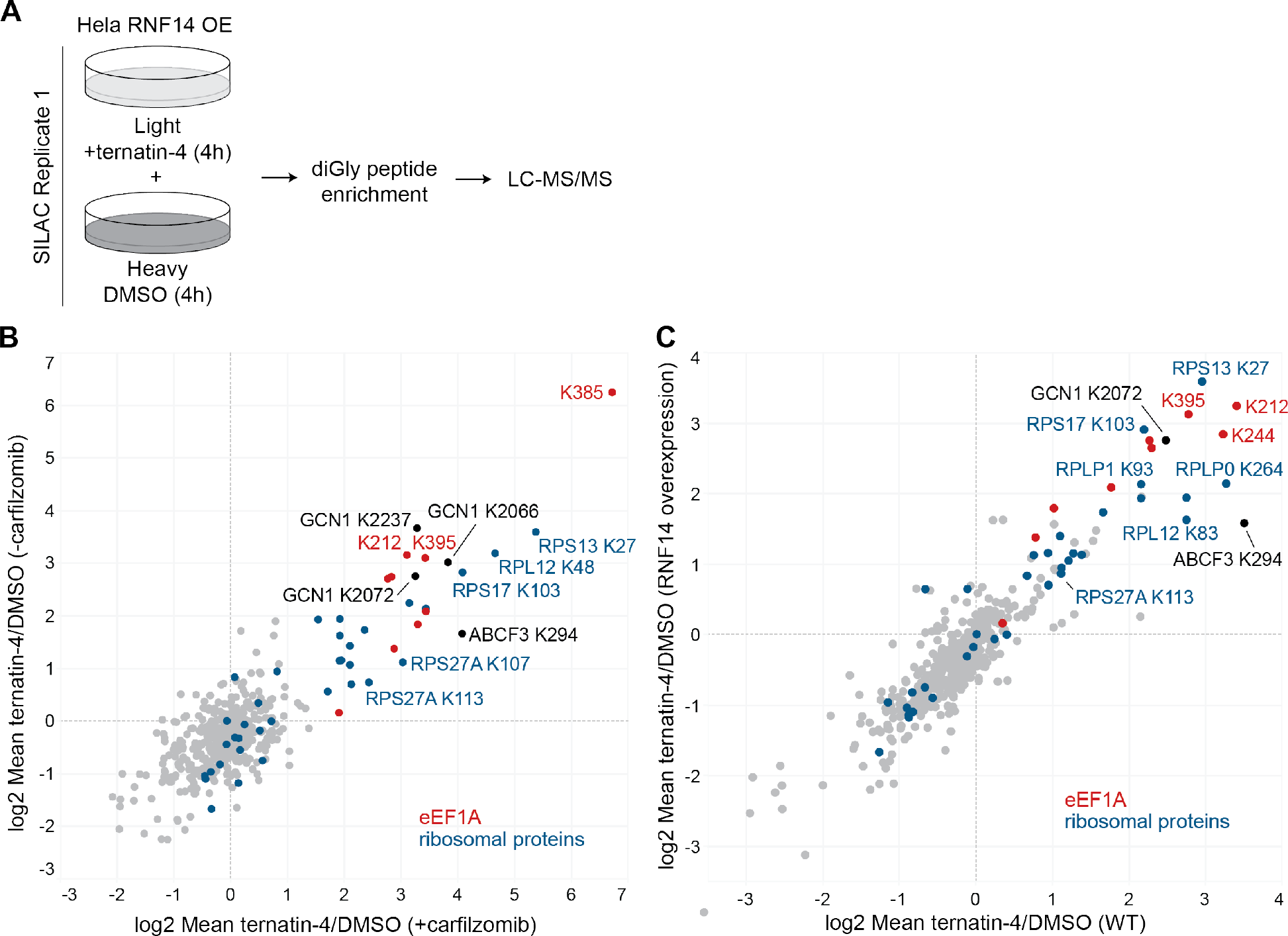
Ternatin-4 promotes RNF14 and RNF25-dependent ubiquitination of eEF1A and ribosomal proteins. Related to. Figure 4. (**A**) Schematic for the SILAC-based diGly proteomics method to identify ternatin-induced ubiquitination sites. Sample configuration is shown for the experiment in Figures 4A and 4B. Replicate 2 was performed identically except heavy and light labels (Arg/Lys in growth media) were swapped. (**B**) HeLa cells stably overexpressing RNF14 were labeled in SILAC media as in (**A**) and treated for 4 h with 500 nM carfilzomib ± ternatin-4 (50 nM). Mean SILAC ratios (log2) for each ubiquitination site (2 replicates, label swaps) were plotted against corresponding log2 ratios from the experiment shown in Figure 4A and **4B** (without carfilzomib). (**C**) HeLa cells (without RNF14 overexpression) were labeled in SILAC media as in (**A**) and treated for 4 h ± ternatin-4 (50 nM). Mean SILAC ratios (log2) for each ubiquitination site (2 replicates, label swaps) were plotted against the corresponding log2 ratios from the experiment shown in Figure 4A and **4B** (HeLa RNF14 OE).

**Figure S4.**
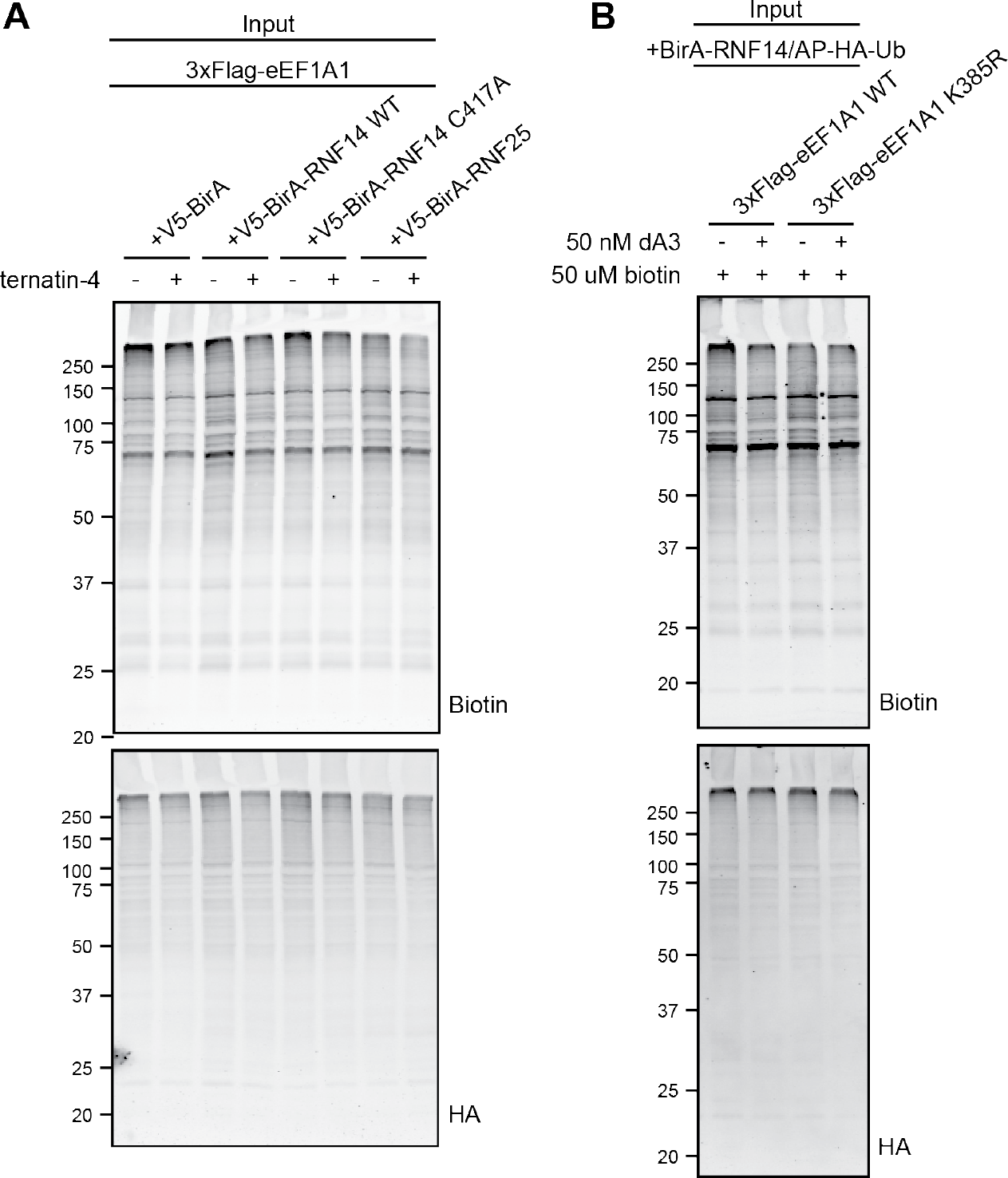
eEF1A K385 is required for efficient degradation and is directly ubiquitinated by RNF14. Related to. Figure 5. Cell lysates from Figure 5D (**A**) and **5E** (**B**) were analyzed by immunoblotting for AP-HA-Ub (anti-HA) and biotinylated proteins (streptavidin).

**Figure S5.**
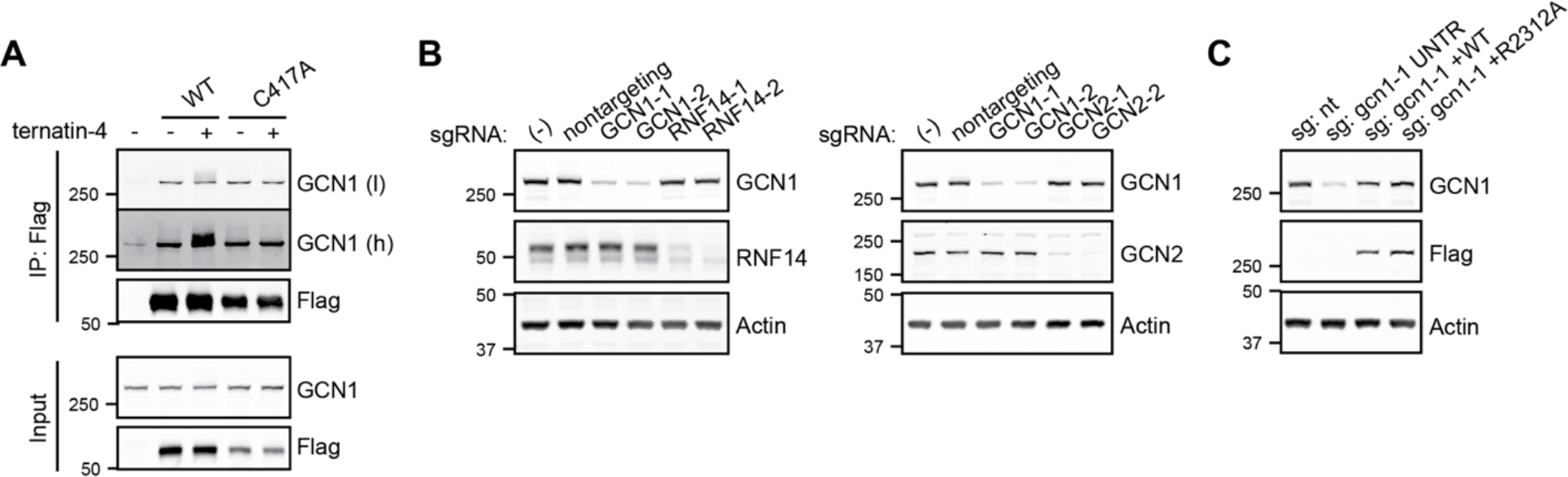
GCN1 interacts with RNF14 and is essential for ternatin-induced eEF1A degradation. Related to. Figure 6. (**A**) Ternatin-induced GCN1 modification requires the catalytic cysteine (C417) of RNF14. As in Figure 6A, HeLa WT cells stably expressing WT or C417A 3xFlag-RNF14 were treated for 4 h with ternatin-4 (50 nM) prior to crosslinking with 0.1% PFA, Flag immunoprecipitation, and immunoblotting. Low and high intensity immunoblot images for GCN1 are shown to indicate higher-MW, presumably multi-ubiquitinated forms of GCN1. (**B**) Immunoblot analysis of lysates from sgRNA-expressing cells used in Figure 6C. Cells were puromycin selected and collected 8 d after sgRNA transduction. All samples were prepared from the same experiment but are depicted on different blots. (**C**) WT and R2312A GCN1 rescue constructs are expressed at similar levels. Cells puromycin-selected for sgRNA expression were transfected with the indicated GCN1-3xFlag_IRES-iRFP constructs to give ∼20% iRFP+ cells. Lysates from the untreated cells used in Figure 6D were analyzed by immunoblotting to show knockdown of endogenous GCN1 and overexpression of exogenous WT and R2312A GCN1-3xFlag.

**Figure S6.**
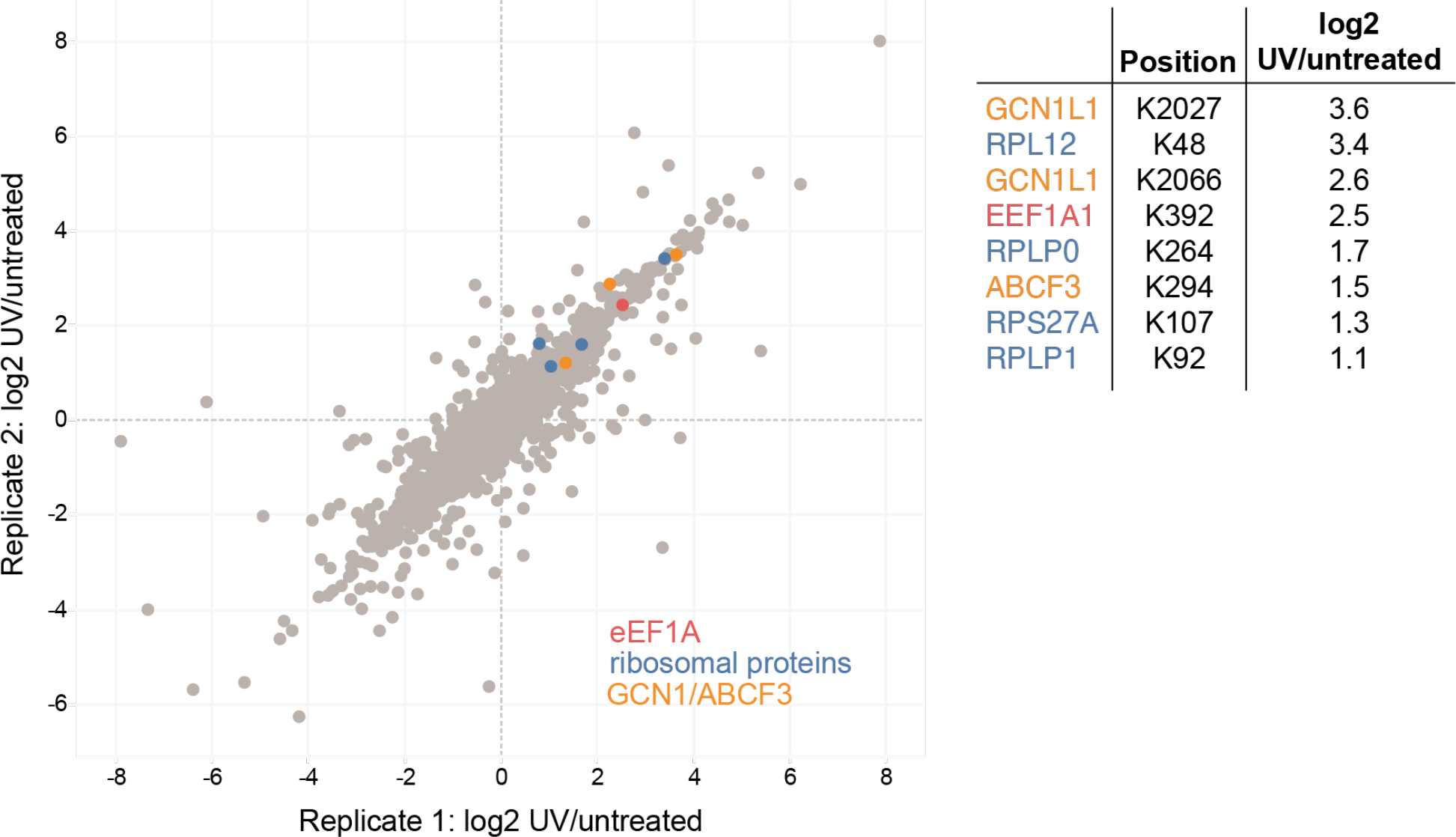
Ternatin-induced ubiquitination sites are also induced by UV irradiation based on previously published data (Elia et al., 2015). Related to Figure 7 and Discussion. Published SILAC diGly proteomics data (Elia et al., 2015) derived from HeLa cells irradiated with UV light (40 J/m^2^) and then allowed to recover for 1 h. Log2 SILAC ratios (UV/untreated) for quantified diGly sites were plotted for two biological replicates. UV-induced diGly sites (log2 SILAC ratio >1) which were also induced by ternatin-4 (Figure 4), are highlighted and log2 SILAC ratios for each site are shown in the table (UV/untreated, mean of 3 replicates).

Supplementary Table 1. CRISPRi screen results, related to Figure 2.

Log2 phenotype score, p value, and gene score (defined as the product of the log2 phenotype score and -log10 p value) for each transcription start site (TSS) targeted in the CRISPRi screen.

Supplementary Table 2. DiGly-modified sites identified by SILAC-MS, related to Figure 4.

DiGly-modified sites and log2 SILAC ratios.

